# The genetic and physiological basis of *Arabidopsis thaliana* tolerance to *Pseudomonas viridiflava*

**DOI:** 10.1101/2023.03.18.533268

**Authors:** Alejandra Duque-Jaramillo, Nina Ulmer, Saleh Alseekh, Ilja Bezrukov, Alisdair R. Fernie, Aleksandra Skirycz, Talia L. Karasov, Detlef Weigel

## Abstract

- The opportunistic pathogen *Pseudomonas viridiflava* colonizes more than fifty agricultural crop species and is the most common *Pseudomonas* in the phyllosphere of European *Arabidopsis thaliana* populations. Belonging to the *P. syringae* complex, it is genetically and phenotypically distinct from well-characterized *P. syringae sensu stricto*. Despite its prevalence, we lack knowledge of how *A. thaliana* responds to its native isolates at the molecular level. Here, we characterize the host response in an *A. thaliana* - *P. viridiflava* pathosystem.
- We measured host and pathogen growth in axenic infections, and used immune mutants, transcriptomics, and metabolomics to determine defense pathways influencing susceptibility to *P. viridiflava* infection.
- Infection with *P. viridiflava* increased jasmonic acid (JA) levels and the expression of ethylene defense pathway marker genes. The immune response in a susceptible host accession was delayed compared to a tolerant one. Mechanical injury rescued susceptibility, consistent with an involvement of JA.
- The JA/ethylene pathway is important for suppression of *P. viridiflava*, yet suppression capacity varies between accessions. Our results shed light on how *A. thaliana* can suppress the ever-present *P. viridiflava*, but further studies are needed to understand how *P. viridiflava* evades this suppression to spread broadly across *A. thaliana* populations.

## INTRODUCTION

Pseudomonads are ubiquitous bacteria of the phyllosphere, the above-ground portion of plants whose main component are the leaves (Vorholt, 2012; Almario *et al*., 2022). There, *Pseudomonas* can play a wide range of roles, from serious pathogens to biocontrol agents (Hirano and Upper, 2000; Legein *et al*., 2020).

*Pseudomonas viridiflav*a is a widespread opportunistic pathogen of plants (Wilkie, Dye and Watson, 1973; Goss, Kreitman and Bergelson, 2005; Lundberg *et al*., 2021), and it is found in a variety of environments, such as leaf litter, rain, and snow (Bartoli *et al*., 2014). Its effects range from no symptoms to growth impairment and disease in soil-grown plants (Jakob *et al*., 2002; Shalev *et al*., 2022), and in axenic conditions it can be lethal for plants (Karasov *et al*., 2018). *Pseudomonas viridiflava* is a natural pathogen of the model plant *Arabidopsis thaliana*, and it has been isolated from *A. thaliana* populations globally (Jakob *et al*., 2002; Goss, Kreitman and Bergelson, 2005; Goss and Bergelson, 2007; Lundberg *et al*., 2021; Karasov *et al*., 2022). In a recent survey of *Pseudomonas* in natural *A. thaliana* populations in Southwest Germany, we found *P. viridiflava* to be the most prevalent *Pseudomonas* clade (Karasov *et al*., 2018). Due to its prevalence and potential to cause disease, *P. viridiflava* likely constitutes a major selective pressure for *A. thaliana* populations (Goss and Bergelson, 2007).

Beyond its interactions with wild plant species, *P. viridiflava* also causes disease in at least fifty different agricultural hosts (Lipps and Samac, 2022). Despite its prevalence and impact in wild and agricultural plant populations (Jakob *et al*., 2002; Goss, Kreitman and Bergelson, 2005; Karasov *et al*., 2018, 2022; Lundberg *et al*., 2021), *P. viridiflava* remains a poorly characterized microbial species. Instead, most of our knowledge regarding *A. thaliana* and *Pseudomonas* interactions comes from *P. syringae sensu stricto* isolates (not *P. viridiflava* isolates), particularly from the model pathogen *P. syringae* pv. *tomato* DC3000 (henceforth Pst DC3000). Originally isolated from tomato, Pst DC3000 is not a natural pathogen of *A. thaliana*, and therefore provides limited insight into the evolution of the *A. thaliana* immune system with its pathogens. The scientific community has nonetheless focused on Pst DC3000 because it easily infects *A. thaliana*, making it very useful for elucidating many of the molecular and biochemical aspects of plant responses to bacterial pathogens (Xin and He, 2013). Typical for many plant parasites, Pst DC3000 uses effector proteins and toxins to manipulate host defense responses and promote bacterial growth. Type III secretion system (T3SS) effector proteins are translocated via the bacterial type III secretion system into the host cells, where they act to suppress host immunity and/or promote disease (Jones and Dangl, 2006; Xin, Kvitko and He, 2018). Several Pst DC3000 effectors can in turn be detected by plant immune receptors, i.e. nucleotide-binding domain leucine-rich repeat (NLR) proteins, eliciting an effector-triggered immune response (Jones and Dangl, 2006). NLR-mediated recognition of effectors activates the salicylic acid (SA) defense pathway to achieve resistance (Glazebrook, 2005). The SA signaling pathway antagonizes the jasmonic acid (JA) signaling pathway (Glazebrook, 2005; Li *et al*., 2019), a property exploited by Pst DC3000: it produces the toxin coronatine that activates the JA defense pathway, consequently decreasing resistance through downregulation of the SA pathway and allowing for increased bacterial growth (Zheng *et al*., 2012; Xin and He, 2013).

Both Pst DC3000 and *P. viridiflava* belong to the *P. syringae* complex, yet they differ substantially in their genetic make-up (Mansfield *et al*., 2012; Gomila *et al*., 2017). In particular, they have different virulence factors (Lipps and Samac, 2022) and thus, the aforementioned mechanisms characterized for Pst DC3000 and related *P. syringae sensu stricto* isolates are not necessarily generalizable to *P. viridiflava*. Indeed, one study has suggested that the defense response of *A. thaliana* to *P. viridiflava* is mediated mainly by the JA defense signaling pathway, with a minor contribution of the SA pathway (Jakob, Kniskern and Bergelson, 2007). This is in contrast with the defense response against Pst DC3000, where SA plays an important role (Glazebrook, 2005; Zheng *et al*., 2012; Xin, Kvitko and He, 2018). Moreover, we have observed that Pst DC3000 is a more virulent pathogen on soil-grown plants than *P. viridiflava* isolates from Southwest Germany are, while many *P. viridiflav*a isolates are more virulent than Pst DC3000 in axenic conditions (Karasov *et al*., 2018; Lundberg *et al*., 2021). *Pseudomonas viridiflava* isolates often have a reduced T3SS effector repertoire, and many encode only *avrE* (Araki *et al*., 2006; Karasov *et al*., 2018), which contributes to the establishment of an aqueous apoplast by manipulating abscisic acid (ABA) signaling, thus resulting in increased pathogen growth (Xin *et al*., 2015; Hu *et al*., 2022; Roussin-Léveillée *et al*., 2022). In addition, *P. viridiflava* isolates from Southwest Germany lack known genes for synthesis of the toxins coronatine, syringomycin, syringopeptin, mangotoxin, phaseolotoxin and tabtoxin (Karasov *et al*., 2018).

While *P. viridiflava* is likely to be among the most, if not the most abundant bacterial pathogen of *A. thaliana* globally, little is known about the mechanisms of virulence in *P. viridiflava* and the corresponding mechanisms of resistance in *A. thaliana*. The differences in the virulence factor repertoire between *P. viridiflava* and Pst DC3000 already suggest that *P. viridiflava* may elicit different host responses compared to Pst DC3000. It has been proposed that the interaction between *A. thaliana* and *P. viridiflava* isolates from the Midwestern USA is not mediated by gene-for-gene recognition, due to the absence of a clear hypersensitive response and the role of JA in resistance (Goss and Bergelson, 2006; Jakob, Kniskern and Bergelson, 2007). The few studies investigating these interactions have focused on bacterial isolates from Midwest USA (Jakob *et al*., 2002; Goss and Bergelson, 2006, 2007; Jakob, Kniskern and Bergelson, 2007), but due to the great intraspecific variation within *P. viridiflava* (Goss, Kreitman and Bergelson, 2005; Bartoli *et al*., 2014; Karasov *et al*., 2018), we do not yet know whether the virulence mechanisms identified in USA isolates are conserved for other *P. viridiflava*.

Here, we used *P. viridiflava* isolated from Southwest Germany in axenic conditions to (i) test for genotype-by-genotype interactions in this pathosystem, (ii) compare the host immune response with what has been described for USA *P. viridiflava* isolates to identify immune responses common to interactions with all *P. viridiflava* and (iii) contrast *A. thaliana* responses to *P. viridiflava* and to the model pathogen Pst DC3000. Through infection trials with a genetically-diverse set of plants and pathogens we assessed the ability of *P. viridiflava* isolates to infect natural *A. thaliana* accessions, identified accessions either susceptible or resistant to *P. viridiflava* infection, and characterized the plant transcriptome and metabolome response upon infection. We found that *P. viridiflava* infection upregulates the JA/ethylene (ET) defense signaling pathway, extending the results obtained with USA isolates to the most prevalent *A. thaliana* pathogen in Europe. Moreover, we found that *P. viridiflava* and Pst DC3000 activate different branches of the JA/ET pathway, underscoring their different pathogenicity mechanisms. Finally, we posit the differences in susceptibility between two closely-related host accessions are due to elevated basal immunity and earlier establishment of a defense response in the tolerant host.

## MATERIALS AND METHODS

For a more detailed description of the methods and data analysis, see Methods S1.

### Biological material

As hosts, we used 21 natural *Arabidopsis thaliana* accessions from across the species’ range, namely: Aitba-2, Apost-1, Ciste-2, Col-0, Ey15-2, Fei-0, HKT2.4, Jablo-1, Koch-1, Mammo-1, Monte-1, Qui-0, Rovero-1, Sha, Shigu-1, Sij-4, Slavi-1, Star-8, Toufl-1, TueWa1-2, Yeg-1, available as lab stocks (Table S1). F_2_ seeds from a Sha x Sij-4 cross (Chae *et al*., 2014) were used for segregation analysis. As immune mutants, we used *sid2*-2, *eds1*-12, *ein2*-1, *coi1*-16 and *jar1*-1 (Table S2). These mutants are in the Col-0 background, except *coi1*-16 which is in Col-5. *Pseudomonas* isolates were transformed to express the *luxCDABE* operon via electroporation using pUC18-mini-Tn7T-Gm-lux, a gift from Herbert Schweizer (Addgene plasmid # 64963; (Choi and Schweizer, 2006), or via mating with a modified version of it including an origin of transfer (*oriT*).

### Axenic infections

Drip-inoculation of 11 day-old plants with *Pseudomonas viridiflava* or *Pseudomonas syringae* pv. *tomato* DC3000 was performed as described before (Karasov *et al*., 2020).

### Luminescence and green pixels quantification

We quantified luminescence of *luxCDABE*-expressing bacteria as a proxy for *in planta* bacterial growth. Whole rosettes of infected plants were harvested and transferred to 96-deep-well plates (2.2 mL, Axygen), containing two 5 ± 0.03 mm glass beads (Roth) and 400 μL of 10 mM MgSO_4_, and ground for two minute at 20 Hz in a TissueLyser II (QIAGEN). Then, 10 mM MgSO_4_ was added to a final volume of 1 mL, and 200 μL were transferred to a 96-well Lumitrac white plate. Luminescence was measured in a TECAN Spark multiplate reader with 2000 ms of integration time. Each well was measured three times, and the mean was calculated for further analysis. The signal of 10 mM MgSO_4_ blanks was subtracted from the samples’ signal before analysis. A constant value equal to the lowest value was added to all samples before log_10_-transformation, and these log_10_-transformed data were used for comparisons.

### Transcriptomics

Single rosettes were harvested at 0, 4, 16, 42 and 72 hours post infection in 2 mL tubes containing two 5 ± 0.03 mm glass beads (Roth). Samples were snap-frozen in liquid nitrogen and stored at -80ºC until RNA extraction. Plants were ground for 30 seconds at 25 Hz in a TissueLyser II (QIAGEN) and RNA was extracted as described (Yaffe *et al*., 2012) using EconoSpin plate columns (Epoch Life Science). mRNA libraries were prepared following an in-house protocol (rCambiagno *et al*., 2021). Multiplexed libraries were single-end (150 bp) sequenced on a HiSeq3000 instrument (Illumina). Reads were mapped against the *A. thaliana* TAIR10 reference genome with bowtie2 v2.2.3 (Langmead and Salzberg, 2012), and transcript abundance was calculated using RSEM v1.2.31 (Li and Dewey, 2011). Default parameters were chosen unless mentioned otherwise. Differential gene expression analyses were performed in R v4.1.0 (R Core Team, 2020) using the DESeq2 package v1.34.0 (Love, Huber and Anders, 2014).

### Metabolomics

Plant rosettes were harvested two days post infection, before evident symptoms were visible. Between 3 and 5 plants were pooled to reach a sample weight of 30 ± 3 mg. Samples were collected in 1.5 mL tubes containing three steel beads of approx. 4 mm diameter (KGM KU 4.000 G28 1.3505 StrG), snap-frozen in liquid nitrogen, and then ground for 30 seconds at 30 Hz in a TissueLyser II (QIAGEN). A methyl *tert*-butyl ether/methanol (MTBE/MeOH) solvent system was used to separate the hormone-containing fraction, metabolite-containing fraction, and protein/starch/cell wall pellet from the ground plant material (Salem *et al*., 2020). This allowed us to simultaneously measure secondary metabolites and hormones from each sample.

## RESULTS

### Host genotype predicts infection outcome with significant host x pathogen interaction

We previously reported that a single diverse *Pseudomonas* clade, OTU5, was prevalent in natural *Arabidopsis thaliana* populations in Southwest Germany (Karasov *et al*., 2018). This clade, classified as *P. viridiflava* (Karasov *et al*., 2018), was the most abundant pseudomonad in *A. thaliana* populations across Europe (Karasov *et al*., 2022). It was recently renamed Pv ATUE5 (*<u>P</u>. <u>v</u>iridiflava* <u>a</u>round <u>Tue</u>bingen 5) (Karasov *et al*., 2022), referencing the region where it was isolated.

A screen of 20 *P. viridiflava* ATUE5 (Pv ATUE5) isolates on a local *A. thaliana* accession, Ey15-2, showed differences in virulence among isolates (Karasov *et al*., 2018), with most isolates being more or similarly virulent as the model pathogen Pst DC3000. To investigate if indeed most Pv ATUE5 isolates are highly virulent under lab conditions, we extended this screen by further infecting the local *A. thaliana* Ey15-2 with 75 additional genetically-diverse ATUE5 isolates from our collection (Fig. 1).

**Fig. 1.**
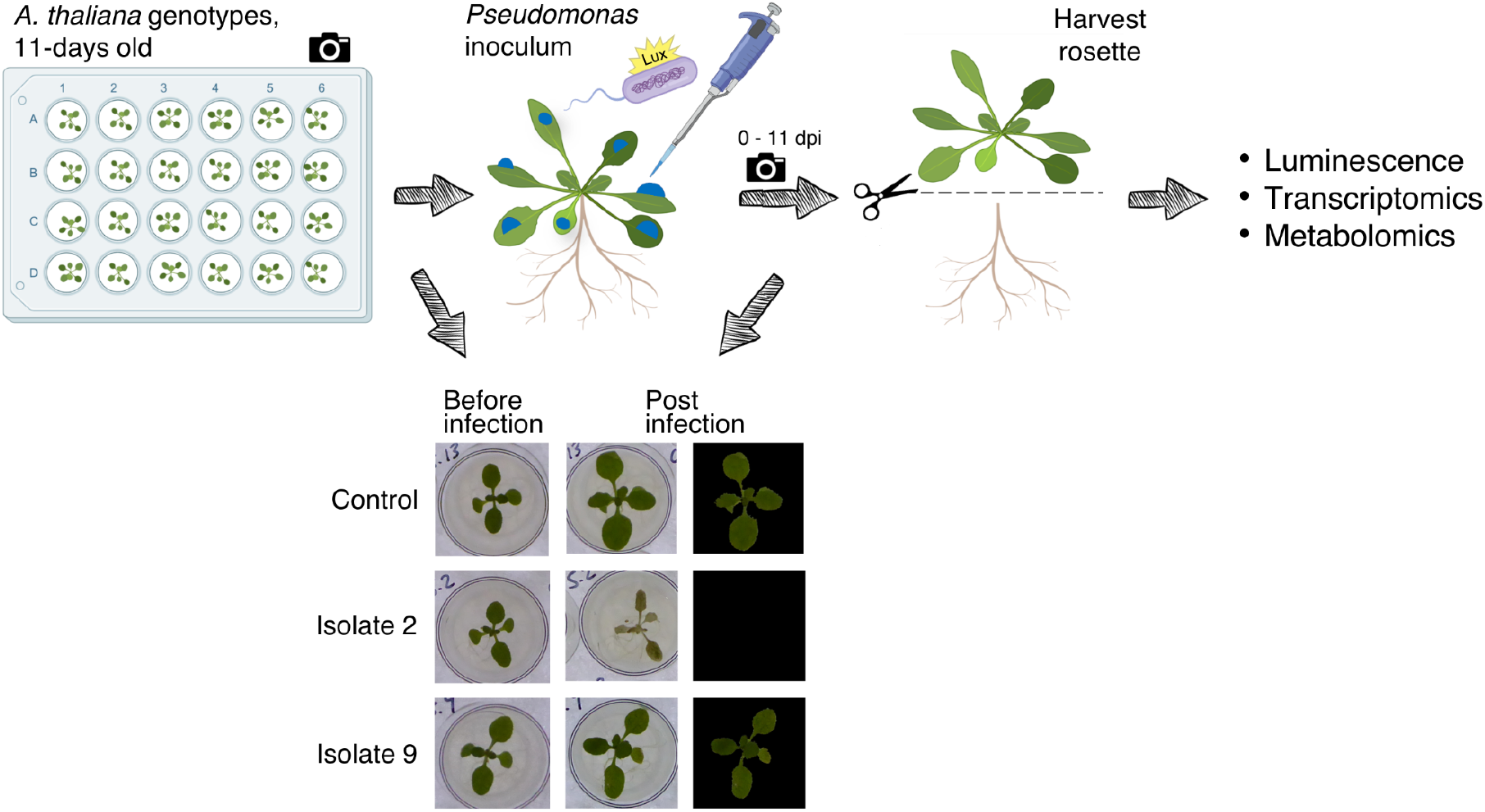
High-throughput phenotyping of infected *Arabidopsis thaliana*. Plants were grown in 24-well plates. When 11 days old, they were drip-inoculated with bacterial inoculum at OD_600_ = 0.01. At selected timepoints, the aerial part of the plant, i.e. the rosette, was harvested for measuring luminescence as a proxy for bacterial growth, for DNA or RNA extraction and sequencing, or for quantification of secondary metabolites and phytohormones. Plates were photographed before infection and/or before harvesting the rosettes in order to quantify the number of green pixels from the images, used as a proxy for non-diseased tissue.

Infection with most of the Pv ATUE5 isolates tested (58/75, 77%) reduced plant growth to the same extent or more than Pst DC3000 (Fig. 2, S1, Table S3). All but one isolate reduced plant size compared to mock-infected plants. This confirms that *P. viridiflava* is a virulent pathogen of *A. thaliana* under axenic conditions. Nonetheless, we observed a large variation in virulence among Pv ATUE5 isolates. This, together with their known genetic variation, could suggest the existence of genotype-by-genotype interactions, where bacterial isolates that are highly virulent on one host accession might be less virulent on others, and vice versa.

**Fig. 2.**
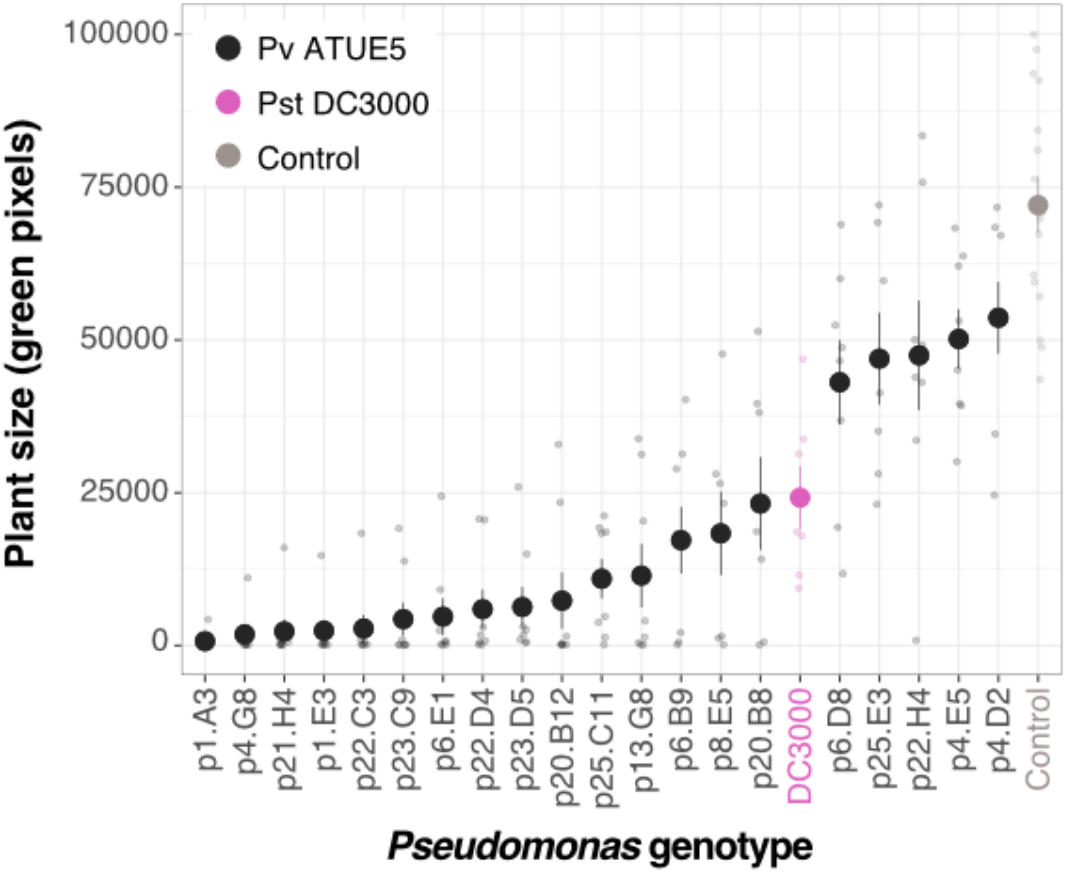
Variation in the virulence of *Pseudomonas viridiflava* ATUE5 isolates on a local *Arabidopsis thaliana* accession. Plant size measured as green pixels 7 days post-infection with the indicated bacterial isolates. DC3000 (pink) corresponds to Pst DC3000, all others (black) are Pv ATUE5 isolates. Small points indicate each replicate, large points indicate the mean, and the whiskers the standard error of the mean. n = 7 -8 for all bacterial pathogens, and n = 16 for the mock-infected plants.

To test whether there are such genotype-by-genotype interactions, we characterized the interaction between a subset of Pv ATUE5 isolates covering the range of virulence (Fig. S1, open circles), and several *A. thaliana* accessions representing the species’ global range and genetic diversity (Table S1, Fig. S2). We drip-inoculated 21 accessions with 12 luminescence-tagged Pv ATUE5 isolates, which allowed us to measure both plant size and bacterial load in the same plants (Fig. 1, Table S4). We confirmed that luminescence signal positively correlated with colony counts (Spearman’s correlation coefficient 0.71, p-value < 2.2×10^−16^), and thus was a valid proxy for bacterial load (Fig. S3a).

Hierarchical clustering of luminescence measurements divided *A. thaliana* accessions into two groups based on pathogen load: either strongly susceptible (high bacterial load) or largely resistant (low bacterial load) to colonization by all Pv ATUE5 isolates tested (Fig. 3a). Sha, Col-0, Yeg-1, Monte-1, Koch-0, Toufl-1, Qui-0, HKT2.4, Rovero-1, Star-8, Mammo-1, and Jablo 1 were susceptible, while Aitba-2, Sij-4, Apost-1, Shigu-1, Ciste-2, Fei-0, TuWa1-2, Ey15-2, and Slavi-1 were resistant. In addition, there were significant differences in the bacterial load between hosts (ANOVA, p-value < 2×10^−16^). We identified Sha as the most susceptible *A. thaliana* accession, with a mean log_10_ luminescence (3.600, SD = 0.541) significantly different from all other accessions (Tukey HSD, p-value < 0.05 for all comparisons; Fig. S4a). Aitba-2 and Sij-4 were the most resistant accessions (mean log_10_ luminescence across all bacterial isolates = 2.312 and 2.464, SD = 0.569 and 0.657, respectively). Despite having very different susceptibility to Pv ATUE5, Sij-4 and Sha were genetically the most-closely related *A. thaliana* accessions included in this study (proportion of SNPs shared = 0.018 based on more than 12 million SNPs; Fig. S2), whereas Aitba-2 was genetically the most distant among all accessions tested (proportion of SNPs shared with other accessions = 0.047 - 0.053), in line with its classification as a relict (1001 Genomes Consortium, 2016). These results suggest that the genotype of the host accession is an important determinant of the outcome of infection, and that even closely-related hosts can differ in their susceptibility to Pv ATUE5.

**Fig. 3.**
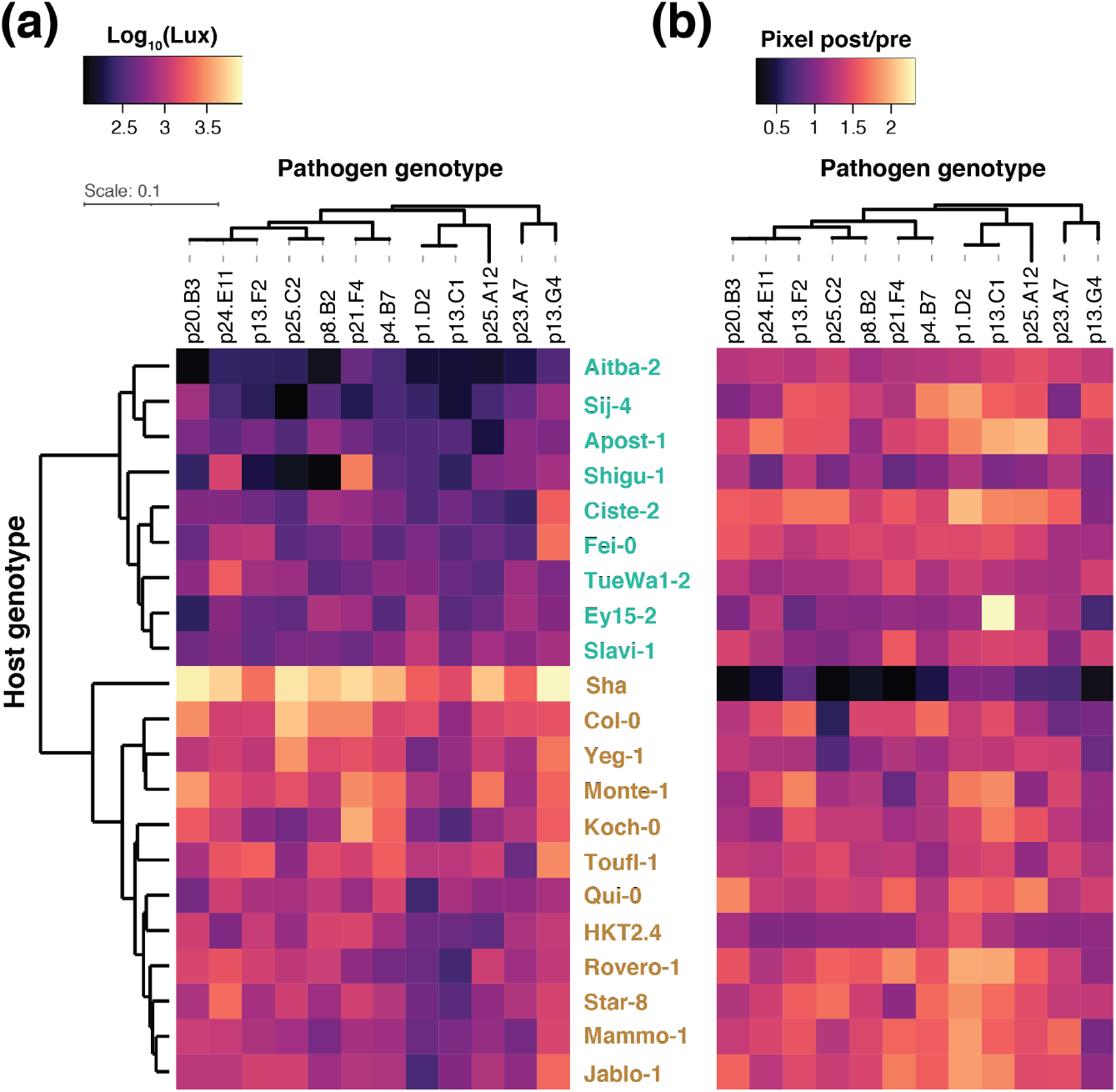
*Pseudomonas viridiflava* ATUE5 load across *Arabidopsis thaliana* accessions. Twenty-one accessions were drip-inoculated with 12 luminescence-labeled pathogen isolates. **(a)** The luminescence signal was measured three days after infection as a proxy for bacterial load. (b) The change in green pixels, expressed as the ratio of measurements before and 3 days after infection, as an indication of plant growth over this interval. Bacterial isolates are ordered based on their phylogenetic relationship, while hosts are ordered by hierarchical clustering based on the luminescence data in **(a)**. Susceptibility based on bacteria load is indicated by the color of the host name: susceptible (ocher) and resistant (aquamarine). n = 8-15 for each host x pathogen combination. We repeated the experiment with a subset of seven host accessions and seven pathogen isolates, which yielded similar results.

Conversely, there were statistically significant differences in the bacterial load each pathogen isolate reached across all host accessions (ANOVA, p-value < 2×10^−16^; Fig. S4b). We identified Pv ATUE5 p13.G4 as the most virulent pathogen (mean log_10_ luminescence = 3.094, SD = 0.660) and isolate Pv ATUE5 p13.C1 as the least virulent one (mean log_10_ luminescence = 2.646, SD = 0.531). Although the range of luminescence across pathogen isolates was narrower than the range across host accessions, bacterial load was significantly different between isolates Pv ATUE5 p13.G4 and Pv ATUE5 p13.C1 (Tukey HSD, p-value < 0.05). The phylogenetic relationships between isolates were not recapitulated by the hierarchical clustering of luminescence signal, with the exception of isolates Pv ATUE5 p13.C1 and Pv ATUE5 p1.D2 (Fig. 3a, S5). Furthermore, we found that closely-related isolates reached different bacterial loads in the same host, supporting the existence of genotype-by-genotype interactions in this pathosystem.

In addition to pathogen load, host-pathogen interactions can be assessed by leaf symptoms, i.e. chlorosis and necrosis, and/or reduced host growth after infection. We used plant growth as an indicator of susceptibility to infection, determined as the change in the number of green pixels estimated from photographs of plants before and after infection (Fig. 1). The change in green pixels during the three days of infection was negatively correlated with bacterial growth as measured by luminescence (Spearman’s correlation coefficient -0.484, p-value < 2.2×10^−16^; Fig. S3b), even though the clustering of host accessions by plant growth was not as obvious as clustering by bacterial load (Fig. 3). This was also the case for the green pixels ratio (Spearman’s correlation coefficient -0.457, p-value < 2.2×10^−16^), and we used the green pixel ratio after and before infection in all experiments where destructive luminescence measurement was incompatible with further sample processing.

Our results so far support the existence of genotype-by-genotype interactions between *A. thaliana* and Pv ATUE5. To quantify these interactions and determine the individual contribution of host and pathogen genotype to infection outcome, we used variance decomposition analysis (Lindeman *et al*., 1980; Grömping, 2015). We found that host genotype explained 16.8% of the variance in luminescence signal, while pathogen genotype explained 3.4%. There was a significant interaction between host and pathogen genotypes (p-value = 0.0076), which explained 7% of the luminescence signal, more than pathogen genotype alone. Fig. S3c shows an example of genotype-by-genotype interactions: Pv ATUE5 p25.C2 reached high load on host Yeg-1, higher than Pv ATUE5 p1.D2 load on the same host. At the same time, the load of Pv ATUE5 p1.D2 was higher than that of Pv ATUE5 p25.C2 in the host Ey15-2. Taken together, these results demonstrate that the extent of bacterial colonization and its impact on host health depend on interactions between pathogen and host genotypes in this pathosystem.

### Colonization progresses differently in susceptible and resistant hosts

The dynamics of *A. thaliana* infection with *P. viridiflava* are unknown. To address this gap in knowledge, we followed the infection of resistant Sij-4 and susceptible Sha hosts over time with two goals: to characterize its progression in terms of bacterial load and host symptom development, and to evaluate if the extent of Sij-4 resistance changed over time. We drip-inoculated Sij-4 and Sha with Pv ATUE5 p13.G4, the most virulent pathogen from our axenic screen (Fig. S4b), and followed infection from 0 to 14 days post-infection (dpi) by both luminescence and green pixels quantification.

We found bacterial load to be highest in the susceptible Sha host at 3 dpi, whereas it took 6 days for bacteria in the resistant Sij-4 host to reach their maximum load (Fig. 4a, Table S5). The maximum bacterial load in both hosts was comparable, yet Sij-4 infected-plants did not lose green pixels despite harboring the same bacterial load associated with tissue collapse in Sha, suggesting that Sij-4 is tolerant to Pv ATUE5 p13.G4. Infected Sij-4 plants had a lower growth rate compared to mock-infected plants, but did not lose green pixels over the course of the experiment (Fig. 4b, Table S6). These results suggest a temporal component in infection outcome, as *P. viridiflava* grows slower in Sij-4, which can additionally withstand a similar bacterial load as the susceptible host with much less impact on plant growth.

**Fig. 4.**
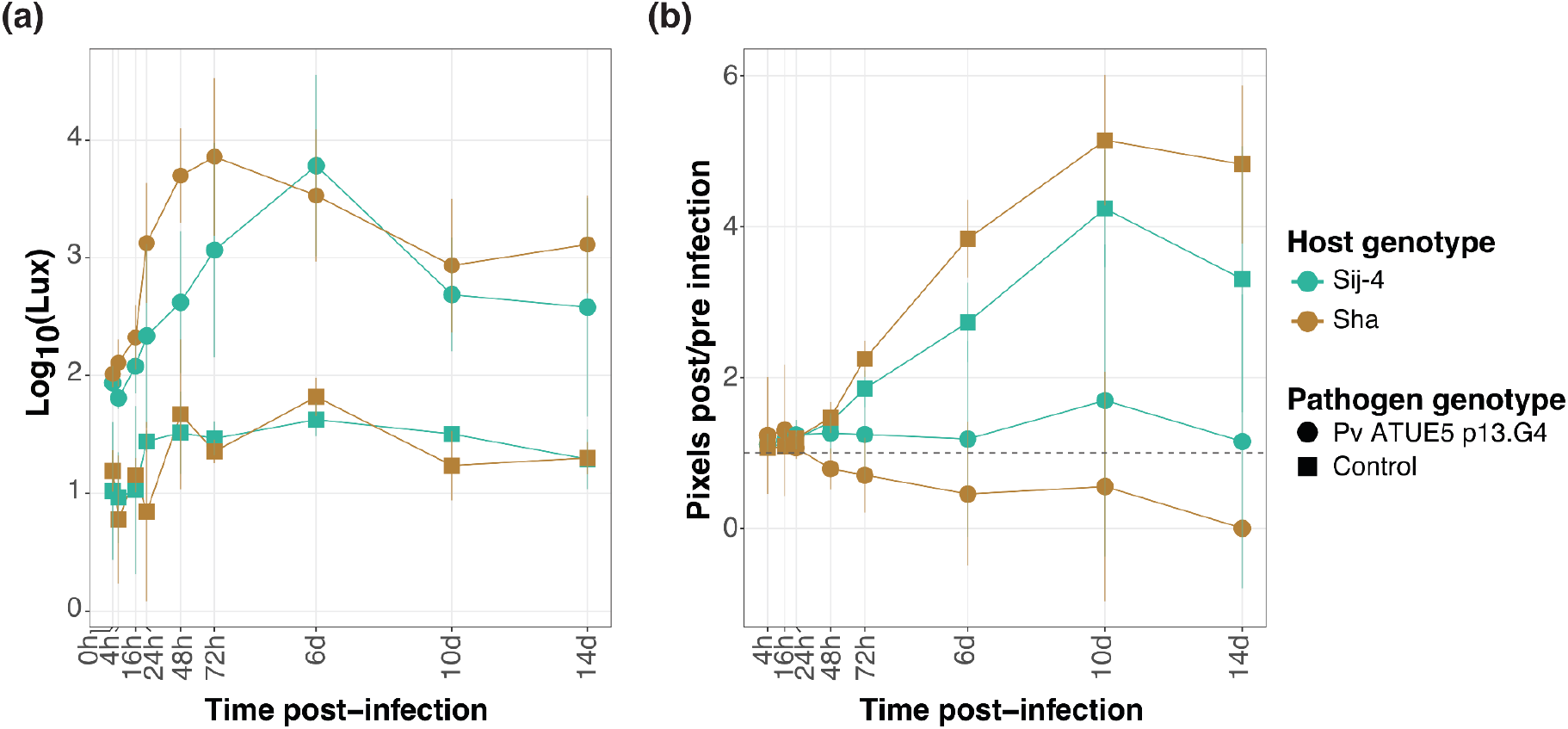
*Pseudomonas viridiflava* growth in different *Arabidopsis thaliana* accessions over time. **(a)** Luminescence over 14 days of Pv ATUE5 p13.G4 infection. **(b)** Mean ratio of green pixels after to before infection (post/pre infection). The dashed line at 1 indicates neither gain nor loss of green pixels. Susceptible Sha plants were all dead after 11 days (0 green pixels ratio). n = 3-6 replicates per host x pathogen x time point for **(a)**; n = 4-24 per host x pathogen x time point for **(b)**.

### Elevated basal immunity in the tolerant host genotype

We next set out to determine the basis of Sij-4 tolerance to Pv ATUE5 infection. First, we analyzed segregation of infection tolerance in an F_2_ population from a cross between tolerant Sij-4 and susceptible Sha (Chae *et al*., 2014). We infected 427 F_2_ individuals and both parental lines in two independent experiments, and measured plant size before infection and 3 days after.

The ratio of green pixels 3 dpi to 0 dpi had a wide distribution among F_2_ plants, indicating that tolerance to *P. viridiflava* might be a polygenic trait (Fig. S6), whereas all mock-infected F_2_ individuals grew similarly. There were no significant size or growth differences between the parental genotypes Sij-4 and Sha when controlling for experiment (pixels before infection ∼ host genotype * experiment, p-value = 0.181; ratio green pixels 3 dpi to 0 dpi ∼ host genotype * experiment, p-value = 0.316). We then estimated the prevalence of tolerance as a trait. For this, we classified F_2_ plants as either more susceptible or more tolerant to Pv ATUE5 p13.G4 based on the ratio of green pixels 3 dpi to 0 dpi of mock-infected F_2_ siblings. The smallest ratio observed in any of the mock-infected F_2_ plants was taken as the threshold for tolerance (experiment 1 = 1.57, experiment 2 = 1.12) (Fig. S6b). Tolerance as defined in this manner was observed in 31% of F_2_ plants in experiment 1 (95% confidence interval [0.257, 0.372]), and 40% in experiment 2 (95% confidence interval [0.331 - 0.473]) (Table 1).

**Table 1.** Prevalence of tolerance of *Pseudomonas viridiflava* ATUE5 p13.G4 infection in a Sha x Sij-4 F_2_ population

| Experiment<br>(threshold) | Host | n tolerant<br>(n total) | Proportion<br>tolerant |
| --- | --- | --- | --- |
| 1<br>(1.57) | Sha x Sij-4 F<sub>2</sub> | 77 (247) | 0.31 |
|  | Sij-4 | 4 (4) | 1.00 |
|  | Sha | 0 (7) | 0.00 |
| 2<br>(1.12) | Sha x Sij-4 F<sub>2</sub> | 72 (180) | 0.40 |
|  | Sij-4 | 12 (14) | 0.86 |
|  | Sha | 2 (10) | 0.20 |

Our results suggest that multiple genes contribute to the tolerant phenotype in Sij-4. While the distribution of green pixels 3 dpi to 0 dpi was largely continuous in the infected F_2_ population, we cannot entirely exclude that there is a single major, recessive locus conferring tolerance. We can, however, exclude that there is a single major, dominant locus conferring resistance, as would be typical for conventional disease resistance conferred by NLR-type *R* genes (Kaur, Bhatia and Mavi, 2021).

### Transcriptome remodeling upon *P. viridiflava* infection

To obtain insight into the potential molecular basis of the differential responses between Sij-4 and Sha to Pv ATUE5 infection, we first compared the leaf transcriptome of the two accessions before infection. There were 270 genes with higher expression in Sha, and 350 with higher expression in Sij-4. A gene ontology (GO) enrichment analysis showed that the top GO terms enriched among genes with higher expression in susceptible Sha were related to iron transport, homeostasis and starvation (Table S7). Meanwhile, the only GO term significantly enriched in tolerant Sij-4 was defense response. These results suggest that the tolerant accession Sij-4 has an elevated basal immunity that could contribute to its tolerant phenotype.

Next, we compared the transcriptional response of Sij-4 and Sha to Pv ATUE5 p13.G4 infection and contrasted it with the well understood response to Pst DC3000 infection. In our axenic system, Pst DC3000 was less pathogenic than *P. viridiflava* (Fig. S7). We collected samples over a 72-hour time course. At the end of the experiment, all Sha plants had a ratio of green pixels 3 dpi to 0 dpi lower than one, i.e. they had lost green pixels, which was not the case for Sij-4 plants.

Principal component (PC) analysis indicated that harvesting time point and infection status contributed the most to the transcriptional profile of the different samples (PC1, 35.9% of variance explained; Fig. S8). The circadian cycle and host genotype had similar effects on the transcriptome: PC2 (14.6% of variance explained) separated samples based on the time of the day/night cycle, while PC3 (13.4% of variance explained) separated samples based on host genotype. These results indicate that the host genotype continues to have a strong effect on gene expression during the course of infection, with drastic transcriptome responses from 42 hpi onwards.

Compared to mock-infected plants collected at the same time point, we observed the largest number of differentially expressed genes (DEGs) in Sha plants infected with Pv ATUE5 p13.G4: over 4,000 genes were up- or down-regulated at 42 hpi, and over 8,000 at 72 hpi (Fig. 5a). The number of DEGs in Pst DC3000-infected Sha plants was much smaller, between one eighth and one fifth of DEGs compared to Pv ATUE5 p13.G4 infection. In Pv ATUE5 p13.G4*-*infected Sij-4 plants, the largest number of DEGs was also observed at 72 hpi: over 2,500 DEGs (Fig. 5b). Most of the DEGs were upregulated, which was true for all host x pathogen combinations.

**Fig. 5.**
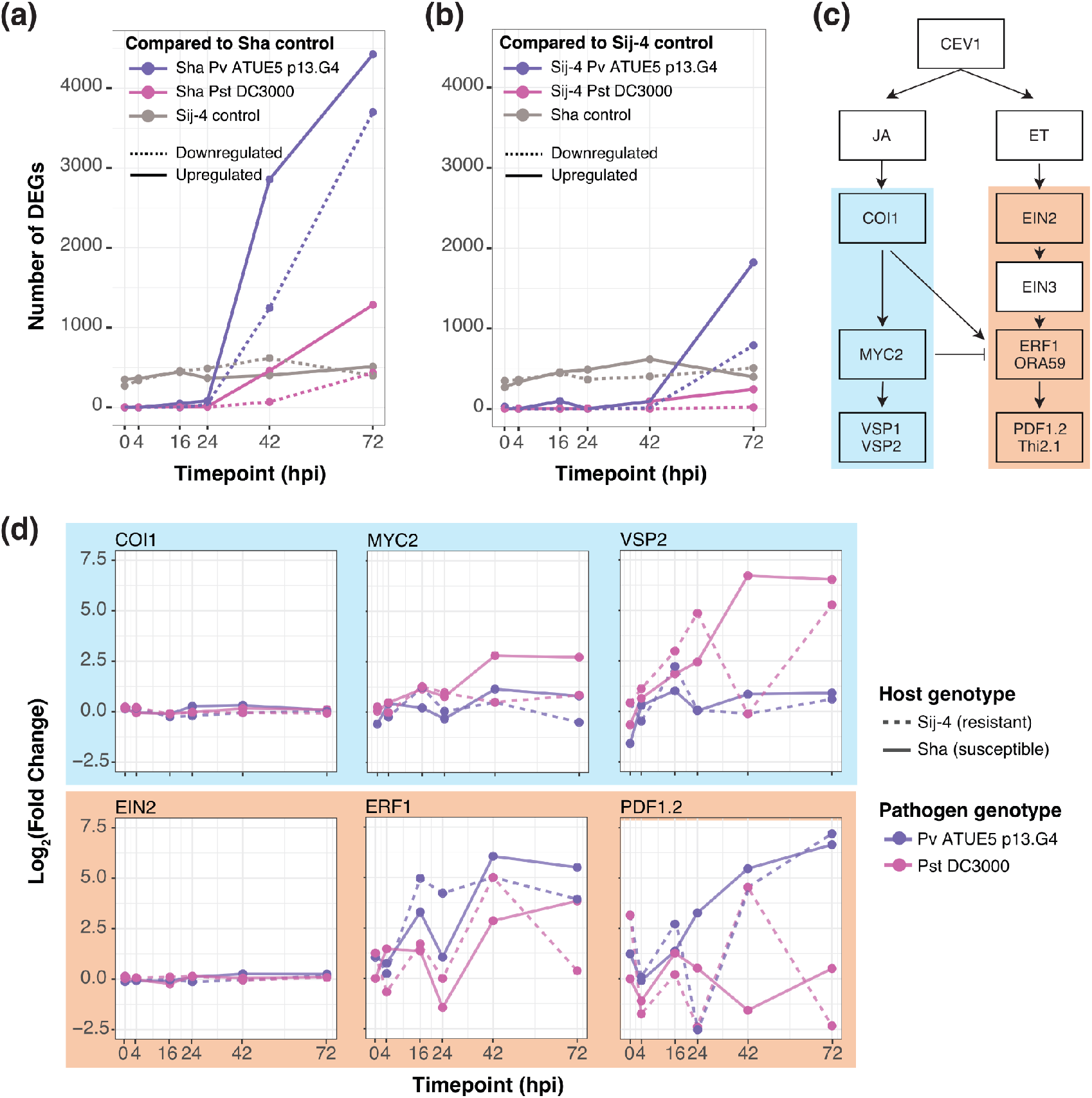
Upregulation of different branches of the JA/ET defense pathway by *Pseudomonas viridiflava* ATUE5 and *Pseudomonas syringae* pv. *tomato* DC3000. **(a)** Number of DEGs over time in infected Sha plants compared to mock-infected plants at the same time point. **(b)** Number of DEGs over time in infected Sij-4 plants compared to mock-infected plants at the same time point. **(c)** Schematic representation of the JA/ET defense signaling pathway. **(d)** Log_2_ fold change of marker genes of the JA/ET defense signaling pathway in tolerant Sij-4 and susceptible Sha hosts relative to mock-infected plants at the same time point. In **(a)** and **(b)**, solid line: upregulated genes, dashed line: downregulated genes; in **(d)**, solid line: Sij-4, dashed line: Sha. Gray: mock-infected, pink: Pst DC3000, purple: Pv ATUE5 p13.G4. n = 3-5 replicates per host x genotype x time point combination, except for Sij-4 x Pst DC3000 × 24 hpi where n = 2. DEGs: differentially expressed genes.

We noted that there were more DEGs in Sij-4 than in Sha infected plants at early time points, particularly at 0 and 16 hpi. An earlier transcriptional response in Sij-4 upon infection compared to Sha is consistent with an elevated basal immunity of Sij-4 and could underlie its tolerant phenotype. To determine whether defense related genes were elevated in Sij-4, we performed a GO enrichment analysis of the DEGs between infected and mock-infected Sij-4 plants at 0 (35 DEGs) and 16 hpi (99 DEGs). Because at 4 hpi there were only 3 DEGs, this time point was not included. While no significant GO term was enriched at 0 hpi, the 98 upregulated genes in infected Sij-4 plants at 16 hpi were enriched in GO terms related to JA as well as immune and defense responses (Table S8). This was not the case for the 49 upregulated genes in infected Sha plants at 16 hpi. Instead, and different from Sij-4, top enriched GO categories in Sha at 16 hpi included detoxification/response to toxic substances and response to hypoxia (Table S8). GO categories enriched in both Sij-4 and Sha after infection compared to control were related to response to biotic stimulus and fungus.

To determine whether Sha responds in the same manner as Sij-4 but with a temporal delay, we extracted the genes belonging to GO categories related to JA and wounding, a known inducer of JA, identified in Sij-4 at 16 hpi and analyzed their behavior in Sha. A majority (13 and 9 out of 17) were indeed significantly upregulated in Sha at 42 hpi and 72 hpi, respectively. Taken together, these results indicate that the immune response in both tolerant Sij-4 and susceptible Sha hosts is qualitatively similar, but Sij-4 mounts a JA-mediated immune response to infection by Pv ATUE5 quicker than the more susceptible accession Sha. JA is known to mediate defense against necrotrophic pathogens and herbivory (Glazebrook, 2005; Mengiste, 2012), and a previous study with the Col-0 reference strain of *A. thaliana* has already demonstrated with JA pathway mutants that this hormone is important for an effective defense against *P. viridiflava* (Jakob, Kniskern and Bergelson, 2007).

To analyze the involvement of the JA/ET pathway in more detail, we compared the expression of well-known marker genes *COI1, MYC2, VSP2, EIN2, ERF1* and *PDF1*.*2* upon infection with Pv ATUE5 p13.G4 and with Pst DC3000 (Fig. 5c). Infection with Pv ATUE5 induced *ERF1* and *PDF1*.*2*, while Pst DC3000 infection induced *MYC2* and *VSP2* (Fig. 5d). This differential upregulation of the ET and MYC2 branches of the JA/ET defense pathway was observed in both Sij-4 and Sha hosts. Taken together, these results underscore the role of the JA/ET pathway in defense against *P. viridiflava*, as seen with different isolates of this pathogen and a different *A. thaliana* host accession (Jakob, Kniskern and Bergelson, 2007).

### Increased accumulation of defense-related hormones in response to *P. viridiflava* infection

To test whether increased transcription of marker genes for the JA/ET signaling pathway reflected increased biosynthesis of JA, we measured the abundance of JA, its precursors and its active form JA-Ile upon infection. We drip-inoculated Sij-4 and Sha plants with Pv ATUE5 and Pst DC3000 and harvested plants 2 dpi, before the susceptible Sha displayed disease symptoms.

In both Sij-4 and Sha plants, Pv ATUE5 infection induced JA and JA-Ile as well as several JA precursors compared to mock- and Pst DC3000-infected plants (Dunn’s test, adj. p-value < 0.05; Fig. 6, Tables S5, S6). The last intermediates before dedicated JA synthesis, OPC-4/-6/-8, were detected almost exclusively in Pv ATUE5-infected plants (Fig 6b). Between the two accessions, the JA precursor FA 18:3 was already significantly more abundant in mock-infected Sha than in Sij-4 plants (Kruskall-Wallis, p-value = 0.028). The differences increased upon Pv ATUE5 infection, with JA and JA-Ile as well as all precursors except OPDA and dinorOPDA being induced to a greater extent in Sha than in Sij-4 plants (Kruskal-Wallis, p-value < 0.05).

**Fig. 6.**
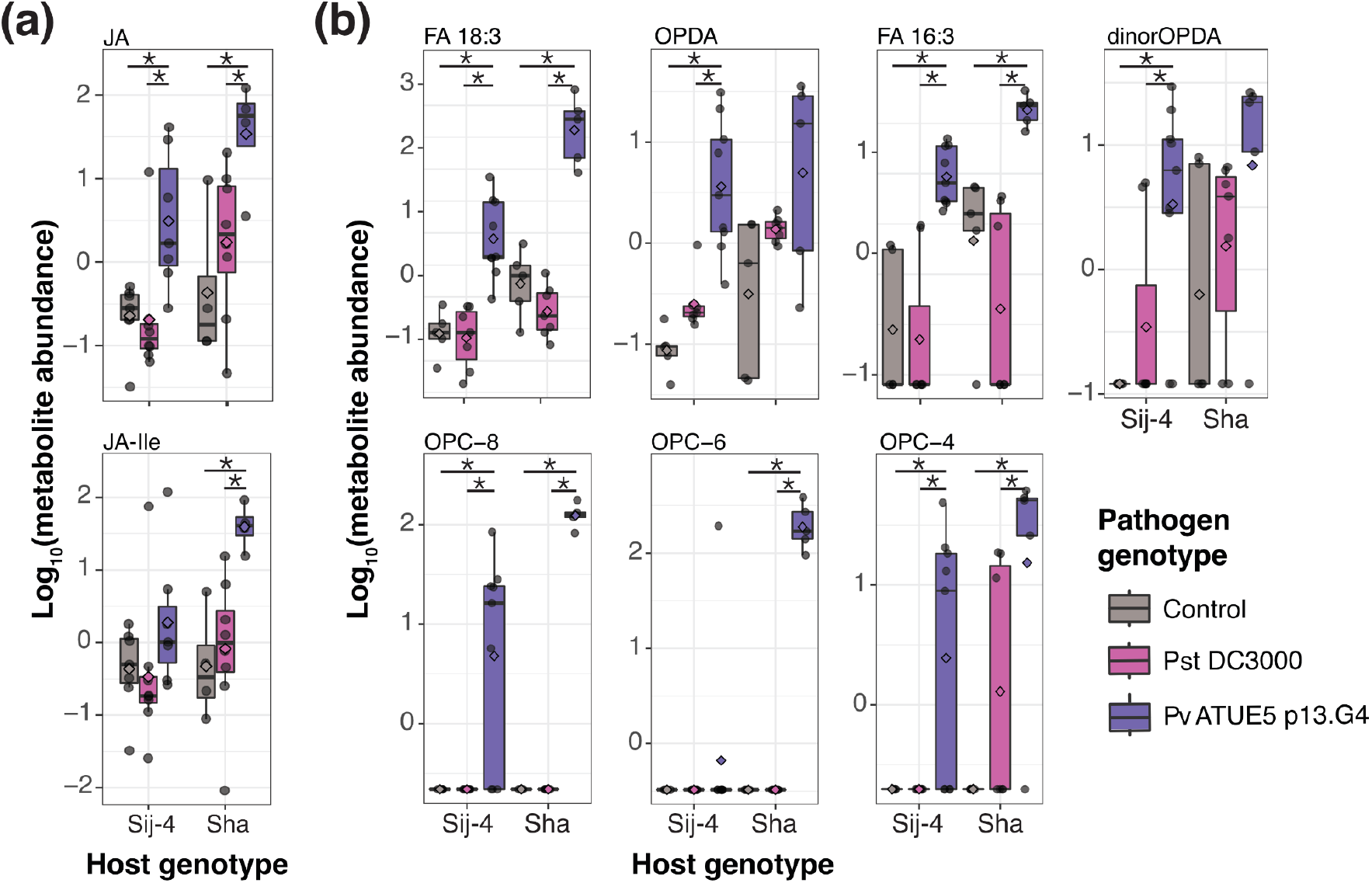
*Pseudomonas viridiflava* ATUE5 infection increases the abundance of JA. **(a)** Response of JA and JA-Ile to *Pseudomonas* infection. **(b)** Response of JA biosynthesis intermediates to *Pseudomonas* infection. JA: jasmonic acid, JA-Ile: JA-isoleucine conjugate, FA: fatty acid, OPDA: oxophytodienoic acid, OPC-8: oxo-pentenyl-cyclopentane octanoic acid, OPC-6: oxo-pentenyl-cyclopentane hexanoic acid, OPC-4: oxo-pentenyl-cyclopentane butanoic acid. Gray: mock-infected control, pink: Pst DC3000, purple: Pv ATUE5 p13.G4. n = 4-9 replicates per host x pathogen genotype combination. Each replicate was a pool of 3 to 5 individual plants. Asterisks indicate statistically significant differences between groups (Dunn’s test, adj. p-value < 0.05). p-values adjusted for multiple comparisons using the Benjamini-Hochberg method.

To determine the influence of Pv ATUE5 infection on other metabolites, both related and unrelated to defense, we performed untargeted metabolomics. In addition to the JA precursors described above, 527 lipids, metabolites and dipeptides varied in abundance among samples (Table S9). Pv ATUE5 lacks genes for the biosynthesis of the toxins coronatine and phevamine, which are known to contribute to *P. syringae* virulence (Zheng *et al*., 2012; Xin and He, 2013; Karasov *et al*., 2018; O’Neill *et al*., 2018), and we detected these metabolites only in Pst DC3000-infected samples (Fig. S9a). The overall metabolomic profile of either Pst DC3000- or Pv ATUE5-infected plants differed considerably from that of mock-infected plants (Fig. S9c). Pv ATUE5 had a greater effect on the host metabolome than Pst DC3000, with the susceptible host Sha having a stronger response to infection than the tolerant host Sij-4.

Host and pathogen made similar contributions to the change in metabolite profile, with the host explaining 18% of the variance (PERMANOVA, p-value = 0.0001) and the pathogen, 22% (p-value = 0.0001). There was a significant genotype-by-genotype interaction, explaining 10% of the secondary metabolomic profile (p-value = 0.098). The distribution of mock-infected samples in the PCoA indicated that, similar to the transcriptome, the metabolome of Sij-4 and Sha already differed before infection (Fig. S9c), which was supported by PERMANOVA (R^2^ = 0.365, p-value = 0.0075). Taken together, genotype-by-genotype interactions influence the outcome of infection not only in terms of the bacterial load but also of the host metabolome, with the most extensive remodeling seen in the susceptible host Sha.

We also measured four other hormones that have been linked in various ways to pathogen defense: salicylic acid (SA), abscisic acid (ABA), and the auxins indole-3-acetic acid (IAA) and indole-3-carboxylic acid (ICA). Similarly to JA and JA-Ile, infection with Pv ATUE5 led to an increase of SA, ABA and ICA, but not IAA, with some differences between susceptible and tolerant hosts; (Fig. 7, Table S10). ABA increased significantly only in susceptible Sha, SA increased significantly only in tolerant Sij-4, and ICA changed similarly in both when compared to mock-infected plants. These differences might contribute to differential host susceptibility to Pv ATUE5 infection. In addition, infection with Pst DC3000 had a smaller impact on hormone levels compared to Pv ATUE5.

**Fig. 7.**
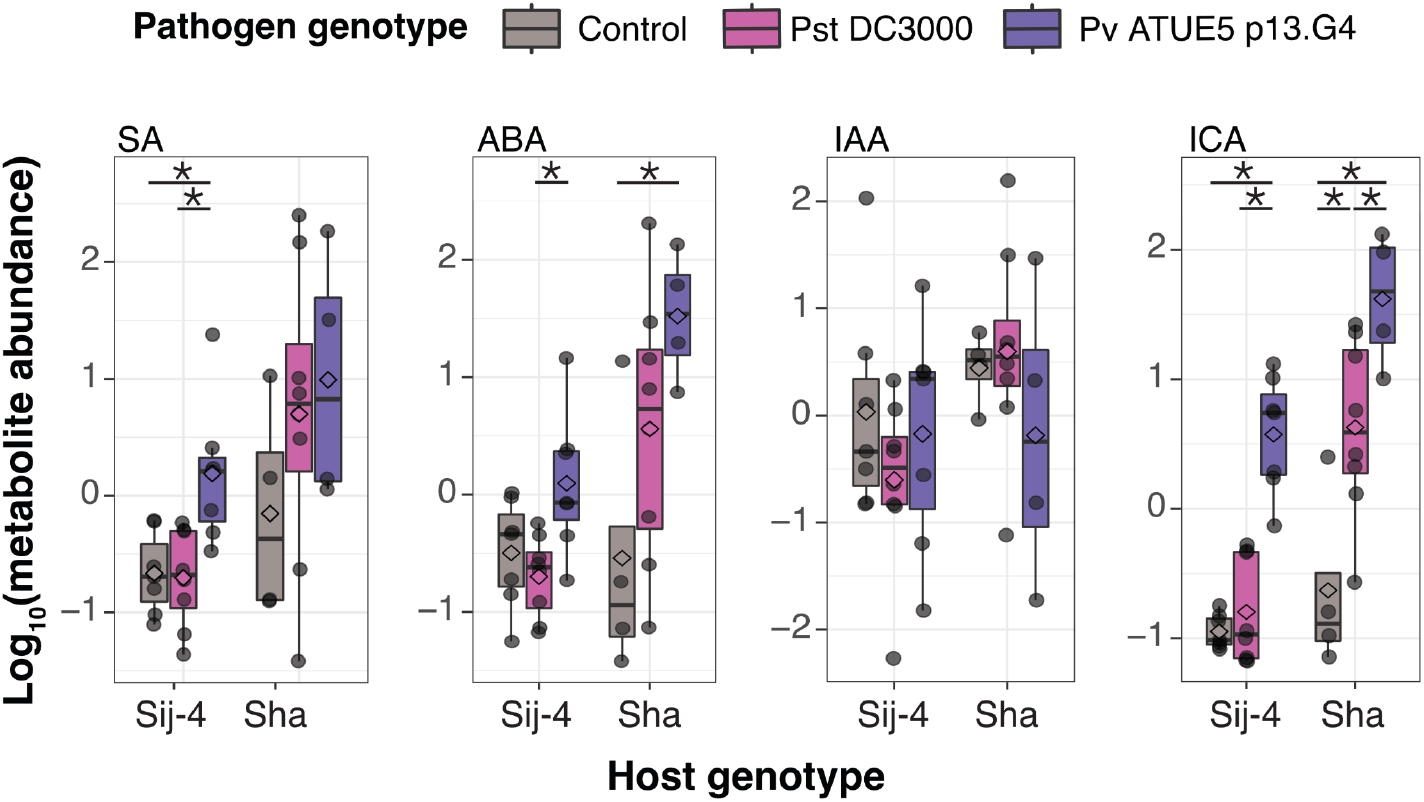
*Pseudomonas viridiflava* ATUE5 infection increases the abundance of several hormones. SA: salicylic acid, ABA: abscisic acid, IAA: indole-3-acetic-acid, ICA: indole-3-carboxylic acid. Gray: mock-infected control, pink: Pst DC3000, purple: Pv ATUE5 p13.G4. n = 4-9 replicates per host x pathogen genotype combination. Each replicate was a pool of 3 to 5 individual plants. Asterisks indicate statistically significant differences between two groups according to Dunn’s test (adj. p-value < 0.05), adjusted for multiple comparisons using the Benjamini-Hochberg method.

### Attenuation of susceptibility to *P. viridiflava* by the JA pathway

Because our transcriptomics and metabolomics results pointed to a role for JA and the ET branch of the JA/ET defense pathway in *A. thaliana* response against Pv ATUE5, we examined Col-0 and Col-5 lines deficient in the SA (*eds1*-12, *sid2*-2), JA (*jar1*-1, *coi1*-16) and ET (*ein2*-1) defense pathways. We found the Col-5 background to be as susceptible to infection as Col-0 (Fig. S10, Table S11), which we used as a control for all mutants.

All SA and JA mutants supported significantly higher Pv ATUE5 load than Col-0 wild type and grew less after infection (Dunn’s test, adj. p-value < 0.05; Fig. 8a,b, Table S11). The most affected mutant was *coi1-*16, and the least affected was *ein2*-1, consistent with our transcriptomics results suggesting JA/ET signaling via COI1 and ERF1, and not EIN2 after Pv ATUE5 infection. Upon Pst DC3000 infection, only the SA mutant *sid2*-2 showed significantly increased bacterial load (Fig. 8a). While not supporting increased bacterial growth like *sid2*-2, *ein2*-1 mutants grew less when infected with Pst DC3000 (Dunn’s test, adj. p-value = 0.003 and 0.025 respectively).

**Fig. 8.**
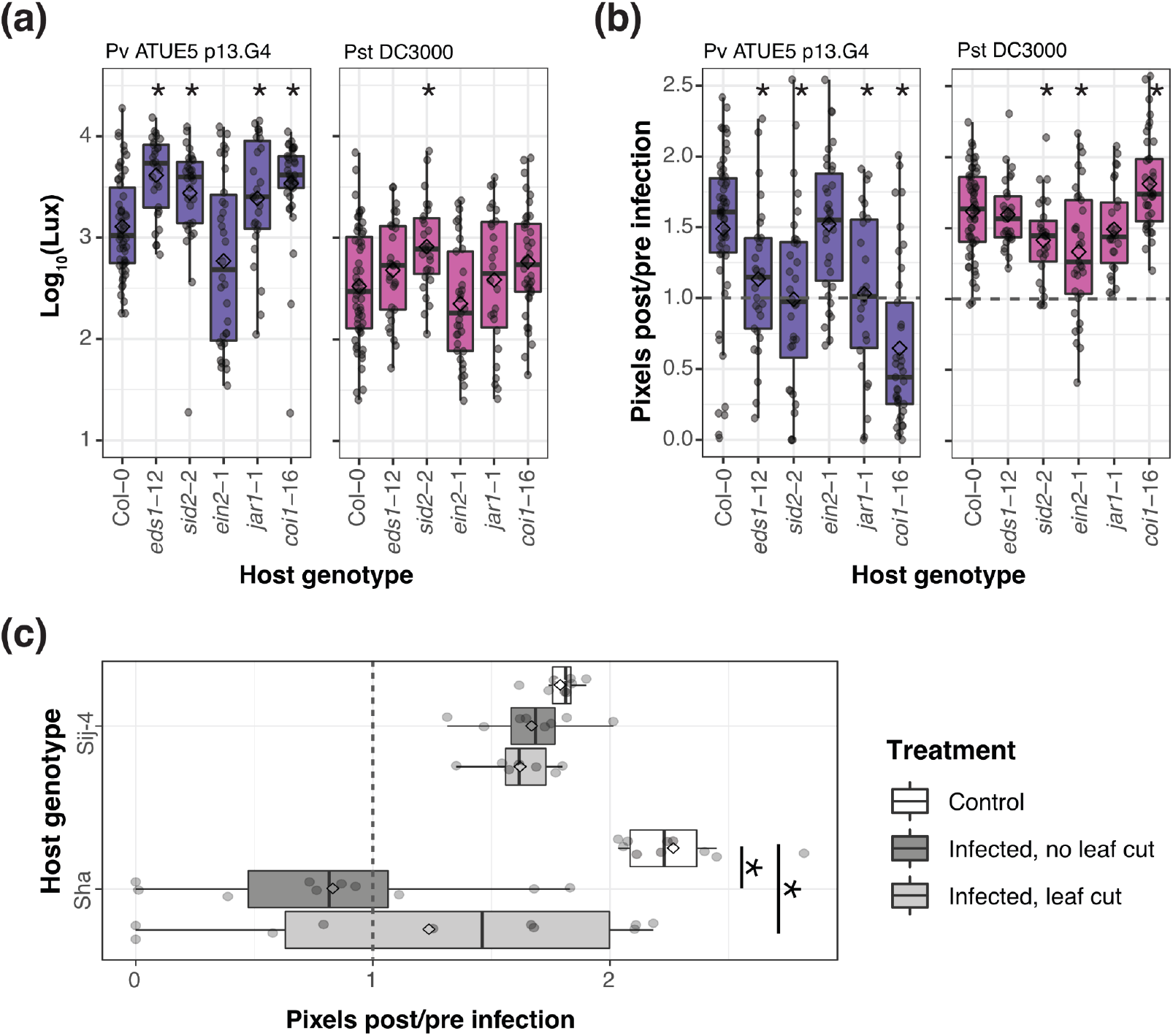
The JA/ET pathway is involved in resistance against *Pseudomonas viridiflava* ATUE5 infection. **(a)** Bacterial load measured as luminescence 3 days post-infection with Pv ATUE5 p13.G4 or Pst DC3000. **(b)** Plant growth measured as the ratio of green pixels 3 days post-infection. Asterisks indicate statistical significance of the Dunn’s test at adj. p-value < 0.05 compared to Col-0 wild type, adjusted for multiple comparisons using the Benjamini-Hochberg method. n = 24-55 plants per host x pathogen genotype combination. Purple: Pv ATUE5 p13.G4, pink: Pst DC3000. **(c)** Plant growth measured as the ratio of green pixels 3 days post-infection to before infection. Infected plants were left intact or had one leaf cut before infection. Asterisks indicate statistically significant differences (two-tailed t-test, adj. p-value < 0.05) between treatments within each host genotype, adjusted for multiple comparisons using the Benjamini-Hochberg method. n = 7-10 plants per host x treatment combination.

Incidentally, we found that Sha susceptibility to Pv ATUE5 p13.G4 decreased when mechanical injury preceded infection. Mechanical injury has been shown to increase JA and its precursors OPDA and dinor-OPDA levels, and to induce changes in defense-related gene expression as early as 15 minutes after injury (Reymond *et al*., 2000). When we cut one leaf of Sij-4 and Sha plants immediately before Pv ATUE5 infection, we observed a decrease in Sha susceptibility compared to non-injured plants (Fig. 8c, Table S12), providing further support for the role of the JA/ET defense pathway in resistance against *P. viridiflava*.

## DISCUSSION

The gammaproteobacterium *Pseudomonas viridiflava*, from the *Pseudomonas syringae* complex, is an opportunistic pathogen that is commonly found in agricultural crops as well as in *Arabidopsis thaliana* populations around the globe (Jakob *et al*., 2002; Araki *et al*., 2006; Bartoli *et al*., 2014; Karasov *et al*., 2018, 2022; Ma *et al*., 2021; Shalev *et al*., 2022). Despite its abundance and likely impact on *A. thaliana* fitness in the wild, little is known about the mechanisms of interaction and pathogenesis in this pathosystem. Understanding the process of infection with *P. viridiflava* is relevant to developing resistance in agriculture as well as to understanding the evolution of the *A. thaliana* immune system. Here, we characterized the interaction between several *A. thaliana* genotypes and closely-related *P. viridiflava* isolates from Southwest Germany, and contrasted the interactions with the response to the well-characterized model pathogen *P. syringae* pv. *tomato* DC3000 (Xin and He, 2013). We demonstrate that tolerance to *P. viridiflava* is mediated by the JA/ET defense signaling pathway, specifically by the ET branch, consistent with experiments using Midwestern USA isolates of *P. viridiflav*a (Jakob, Kniskern and Bergelson, 2007), with greater tolerance in some accessions due to an elevated basal immunity that allows for a faster response to infection.

Past work indicated that the pathogenicity of *P. viridiflava* ATUE5 isolates varies on a single *A. thaliana* genotype (Karasov *et al*., 2018). We have now expanded these results to 75 additional isolates on the local genotype Ey15-2 and to a set of 21 hosts infected with 12 pathogen genotypes. While host genotype was the main determinant of bacterial load, there was a significant genotype-by-genotype interaction, which was also present in the metabolome remodeling of infected plants. Genotype-by-genotype interactions may prevent a single bacterial isolate from taking over in this pathosystem, thus explaining the vast genetic diversity of ATUE5 reported in Southwest Germany (Karasov *et al*., 2018).

We identified the accession Sij-4 as being tolerant to Pv ATUE5 infection. Tolerance is one of the two main strategies of plants successfully coping with pathogens, the other being defense that results in resistance (Pagán and García-Arenal, 2020). Our conclusion was based on measuring both the bacterial load and the effect of infection on plant growth, and it is in line with previous studies reporting tolerance of *A. thaliana* to certain strains of *P. syringae* and *P. viridiflava* (Jakob *et al*., 2002; Kover and Schaal, 2002; Goss and Bergelson, 2006). Genetic determinants of tolerance have been identified in wild and crop plants, ranging from a single or a few large-effect quantitative trait loci (QTL) up to 22 QTL of minor effect (Pagán and García-Arenal, 2020). Tolerance to Pv ATUE5 does not appear to be mediated by a single, dominant locus, as is usually-but not always-the case if it was due to a resistance (R) gene of the NLR class (Dangl, Horvath and Staskawicz, 2013). Reduced activity of negative regulator(s) of the immune defense pathways could underlie the elevated basal immunity of Sij-4 and its faster response to infection compared to the more susceptible Sha accession, resulting in slower pathogen proliferation and less strongly affected plant growth. Indeed, mutations in negative regulators of defense signaling pathways can result in loss-of-susceptibility (Iyer-Pascuzzi and McCouch, 2007; Hashimoto *et al*., 2016), and constitutively primed plants can arise from mutations in several proteins (Mauch-Mani *et al*., 2017), including chloroplastic transcription factors (García-Andrade *et al*., 2011) and MAP kinases (Frye, Tang and Innes, 2001).

A key aspect of tolerance of Pv ATUE5 was the temporal component, with the more tolerant accession Sij-4 slowing pathogen growth and establishing an immune response, mostly linked to JA signaling, as early as 16 hpi. In contrast, a similar JA transcriptional response was observed in the more susceptible Sha accessions only at 42 hpi, apparently too late to control infection. A qualitatively similar but delayed transcriptional response has been observed when mutants defective in multiple defense sectors were challenged with Pst DC3000 and compared to wild-type plants (Mine *et al*., 2018). JA precursors were found to be increased in infected plants regardless of their tolerance to *P. viridiflava*, which might be attributed to both susceptible and tolerant hosts being sampled only two days post-infection, potentially too late to see differences between the two accessions, as the transcriptomes were also more similar at this compared to earlier time points.

Consistent with prior reports of SA being involved in resistance to *P. viridiflava*, although to a lesser extent than JA (Jakob, Kniskern and Bergelson, 2007), SA-deficient mutants were more susceptible to *P. viridiflava* and SA was increased upon *P. viridiflava* infection in the tolerant host, which might contribute to this phenotype. Upon infection with *P. viridiflava*, ABA increased only in the susceptible accession Sha. Pst DC3000 infection can increase ABA levels and ABA signaling, which is associated with stomatal closure and the establishment of an aqueous apoplast, favoring bacterial growth (Hu *et al*., 2022; Roussin-Léveillée *et al*., 2022); and DC3000 manipulation of ABA is mediated by the functionally redundant effectors AvrE and HopM1. AvrE is the only effector in *P. viridiflava* isolates from Southwest Germany (Karasov *et al*., 2018), so *P. viridiflava* could be using similar mechanisms as Pst DC3000 to increase its virulence on susceptible Sha.

We have identified putative tolerance factors on *A. thaliana* against its natural bacterial pathogen *P. viridiflava*, and showed the importance of the JA/ET defense pathway and timing in the defense response of *A. thaliana* to this pathogen. Compared to the model pathogen Pst DC3000, both the transcriptome and metabolome were remodeled more extensively in both Sij-4 and Sha hosts when these were infected with *P. viridiflava*. Consistent with previous reports, we found that Pst DC3000 activates the MYC2 branch of the JA/ET pathway (Zheng *et al*., 2012). Infection with *P. viridiflava*, on the other hand, activated the ET branch of this pathway. Our results suggest that this activation is mediated by an increase in JA and its precursors, which is transduced via COI1 to the ET transcription factor ERF1. Why the activation of the ET branch is achieved via JA instead of ET in this case is not yet clear.

## ACKNOWLEDGEMENTS

We thank Manuela Neumann and Peter Laurie for their help with the axenic screen, Wei Yuan, Max Collenberg and Thanvi Srikant for help with transcriptomics, Jacobo de la Cuesta-Zuluaga for help with statistical analysis, and Andy Gloss for providing luminescence-tagged *Pseudomonas* isolates. This work was supported by an HFSP Long-Term Fellowship (T.L.K.), EU Horizon ERC Synergy Grant 951444 PATHOCOM and the Max Planck Society (D.W.).

## AUTHOR CONTRIBUTIONS

ADJ, TLK and DW devised the study. ADJ, NU and TLK performed the experiments. IB wrote the scripts for image analysis. SA and ARF measured phytohormones. AS measured lipids, metabolites and dipeptides. ADJ analyzed the data. ADJ wrote the first draft of the manuscript with input from all authors. Major edits were provided by TLK and DW.

## DATA AVAILABILITY

Data not available online is available from the corresponding author upon request.

