## Supplemental figures S1-S10 and methods S1 for "The genetic and physiological basis of *Arabidopsis thaliana* tolerance to *Pseudomonas viridiflava*"

### SUPPORTING INFORMATION

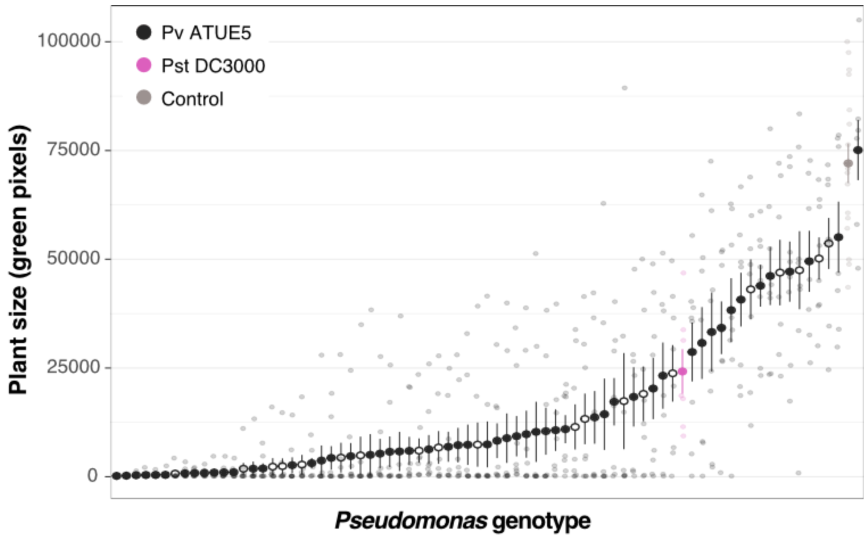


**Fig. S1. Extended screen of Pv ATUE5 isolates virulence on a local *A. thaliana* accession, Ey15-2.** Plant size measured as green pixels 7 days post-infection with the indicated *Pseudomonas* isolates. Points indicate the mean and the error bars indicate the standard error of the mean. *P. viridiflava* isolates shown in Fig. 2 indicated by open circles.

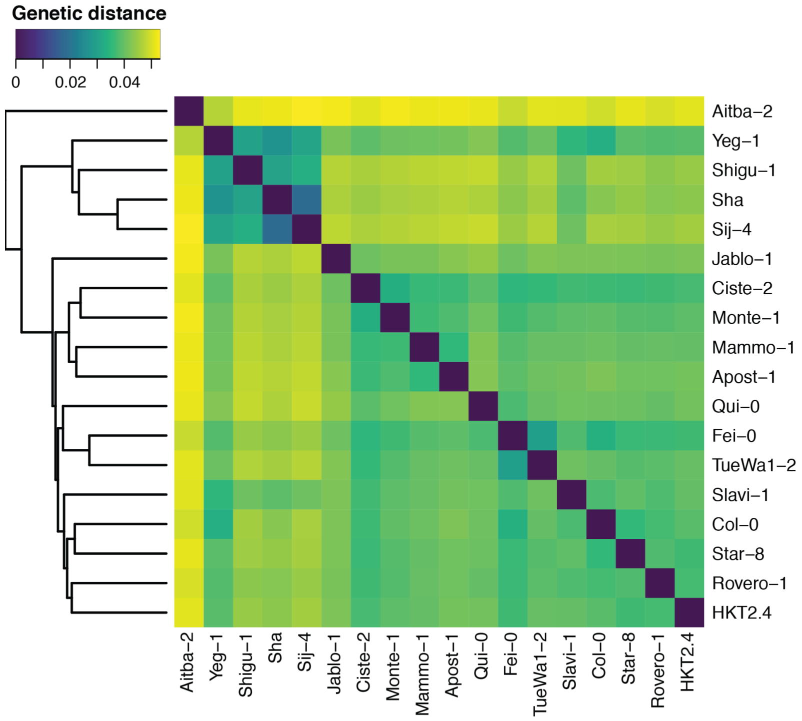


**Fig. S2. Pairwise genetic distance between *A. thaliana* accessions included in this study.** Pairwise genetic distance between 18 *A. thaliana* accessions for which single nucleotide polymorphisms data was available from the 1001 Genomes project (1001Genomes, no date; 1001 Genomes Consortium, 2016). No data was available for Ey15-2, Koch-1 and Toufl-1.

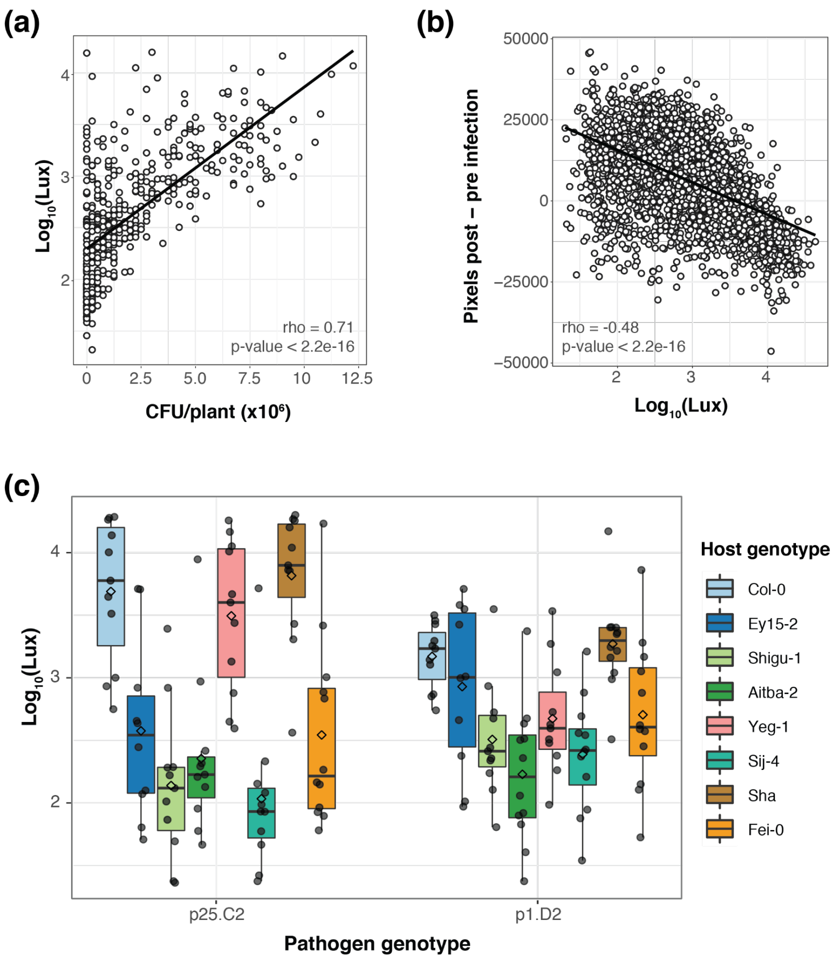


**Fig. S3. Genotype-by-genotype interactions between *A. thaliana* and Pv ATUE5. (a)** Positive correlation between the bacterial load measured as luminescence and the total number of colony forming units (CFU) per plant in a selection of infected plants. **(b)** Negative correlation between luminescence and the ratio of green pixels 3 days post-infection (green pixels 3 days post-infection - green pixels before infection). **(c)** Luminescence 3 days post-infection of two *P. viridiflava* isolates in six selected host accessions. n = 440 plants for A, n = 2,841 plants for B, and n = 8-15 replicates per host x pathogen genotype combination for C.

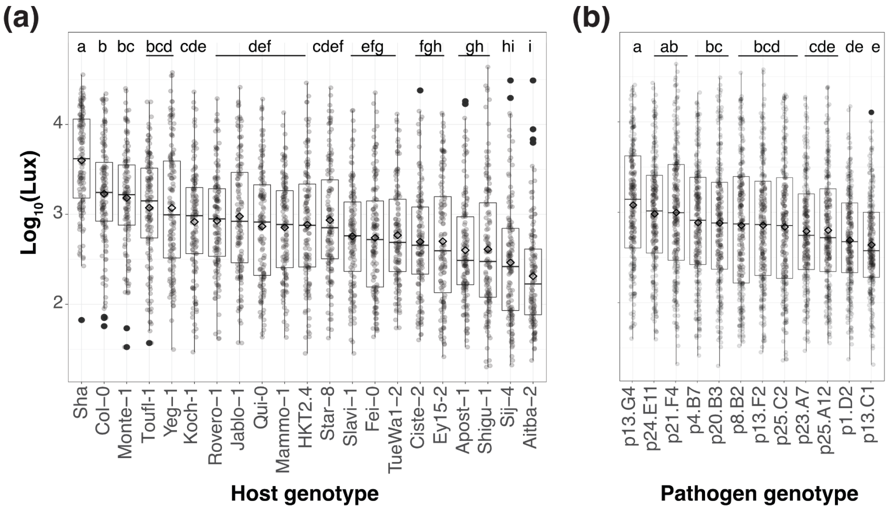


**Fig. S4. Pv ATUE5 growth *in planta* is influenced by host genotype and pathogen genotype to different extent.** Distribution of bacterial load measured as luminescence for all pathogen genotypes on each host genotype **(a)** and of each pathogen genotype across all host genotypes **(b)**. Diamonds represent the mean $log_{10}$(luminescence). Letters above boxplot indicate Tukey's HSD results; genotypes with the same letter were not significantly different from each other at a p-value < 0.05. n = 231-244. We repeated the experiment with a subset of seven host genotypes and seven pathogen genotypes, which yielded similar results.

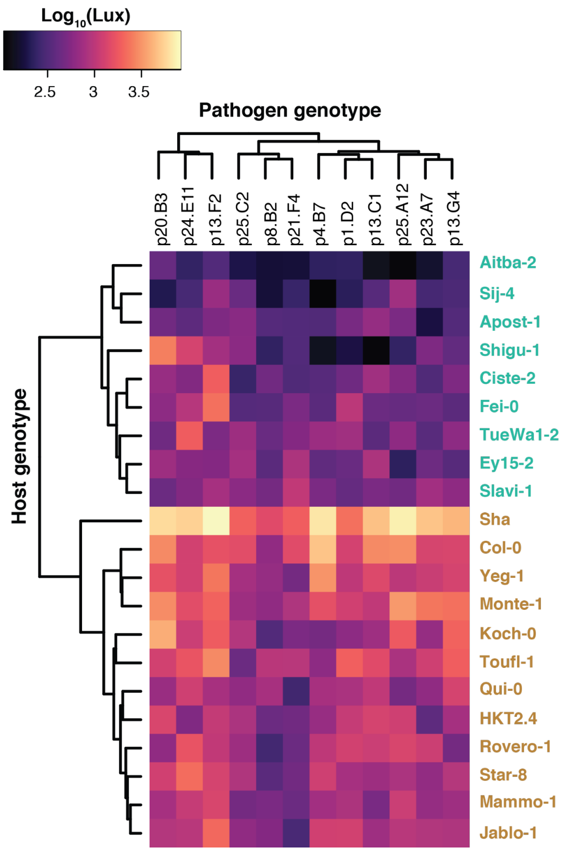


**Fig. S5. Clustering of Pv ATUE5 genotypes based on *in planta* luminescence signal does not recapitulate phylogenetic relationships.** Pathogen load measured as luminescence 3 days post-infection in all host x pathogen genotype combinations. In total, 21 host genotypes were drip-inoculated with 12 luminescence-producing pathogen genotypes. Host and pathogen genotypes are ordered according to hierarchical clustering based on the luminescence data shown in the heatmap. The data presented are the same as in Fig. 3a, but Pv ATUE5 isolates are hierarchically clustered based on luminescence signal instead of their phylogenetic relationship. n = 8-15 for each host x pathogen genotype combination. We repeated the experiment with a subset of seven host genotypes and seven pathogen genotypes, which yielded similar results.

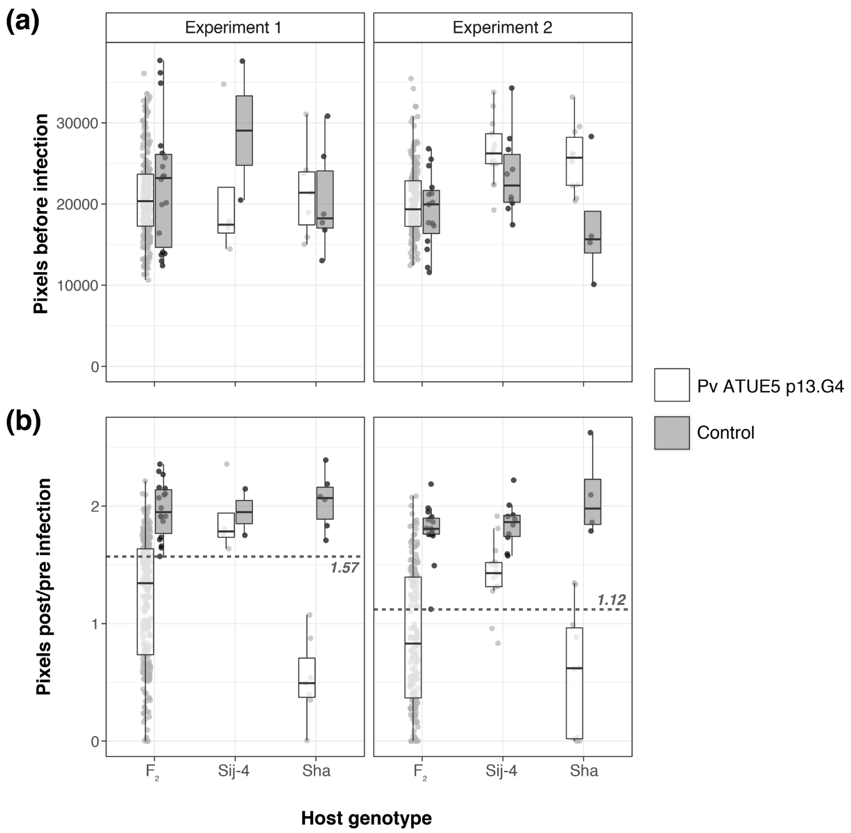


**Fig. S6. Tolerance to Pv ATUE5 is not encoded by a simple dominant locus.** The tolerant Sij-4 accession was crossed to the susceptible Sha accession, and tolerance was scored in the $F_2$ generation in two independent experiments. Green pixels were used as a proxy for plant size. Controls are mock-infected plants of the same genotype (in the case of $F_2$, segregating plants). (a) Plant size before infection. **(b)** Ratio of green pixels 3 days post-infection to before infection. Plants with a ratio > 1 gained green pixels, i.e., were growing, while plants with a ratio < 1 lost green pixels, i.e., were dying. A threshold for classifying plants as susceptible or tolerant was estimated for each experiment based on the ratio of pixels post-/pre-infection of control plants, indicated by the dashed line. Gray: mock-infected plants, white: plants infected with Pv ATUE5 p13.G4.

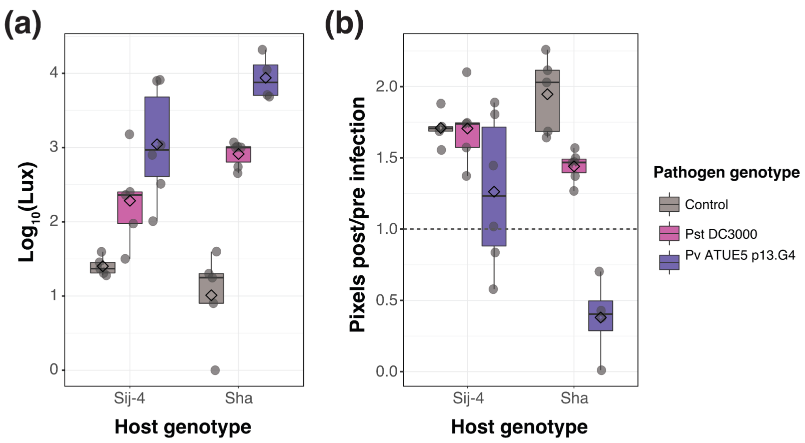


**Fig. S7. *P. viridiflava* is a stronger pathogen of *A. thaliana* under axenic conditions than the model pathogen Pst DC3000. (a)** Bacterial load measured as luminescence 3 days post-infection with Pv ATUE5 p13.G4 (purple) and Pst DC3000 (pink). **(b)** Plant growth measured as the ratio of green pixels 3 days post-/pre-infection in Sij-4 (tolerant) and Sha (susceptible) accessions. The dashed line at 1 indicates neither gain nor loss of green pixels. n = 4-6 plants per host x pathogen genotype combination.

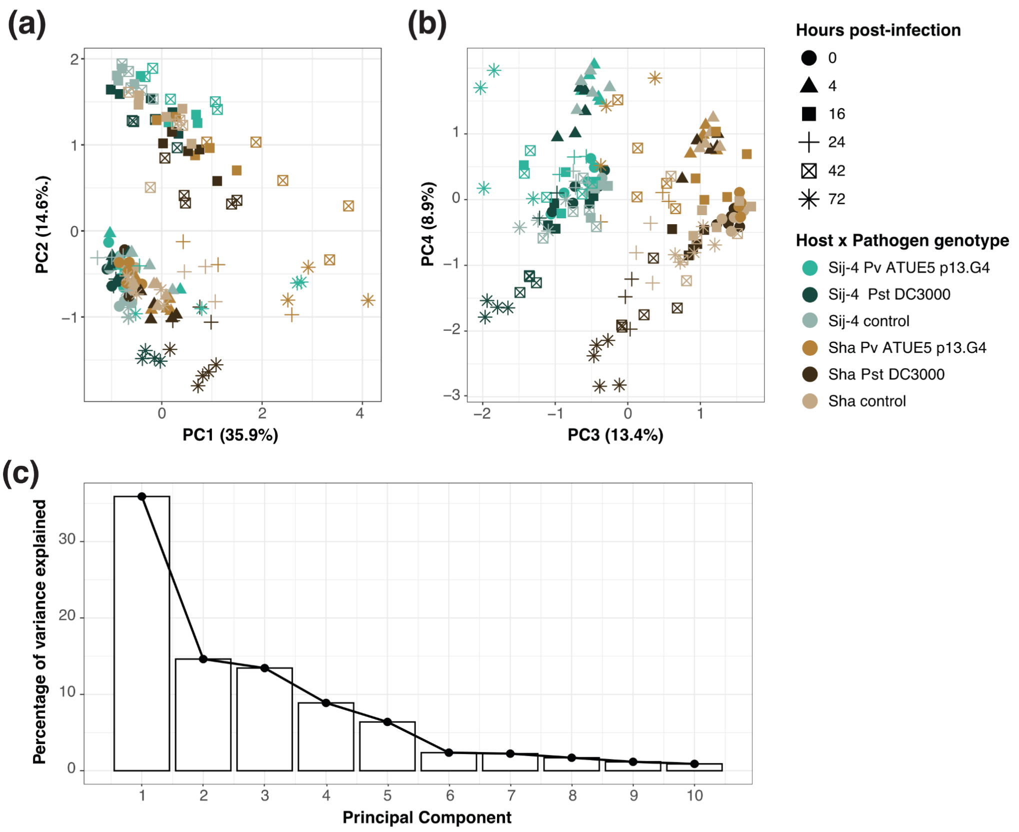


**Figure S8. Principal component analysis of the transcriptome of Sij-4 and Sha infected with Pv ATUE5** p13.G4 **or Pst DC3000.** Plants were harvested at different time points. (a) Principal component 1 (PC1) and PC2. Symbols represent time post-infection at collection, colors represent the host x pathogen genotype combination. (b) PC3 and PC4. (c) Variance explained by the first 10 PCs. n = 3-5 replicates per host x genotype x time point combination, except for Sij-4 x Pst DC3000 x 24 hpi where n = 2.

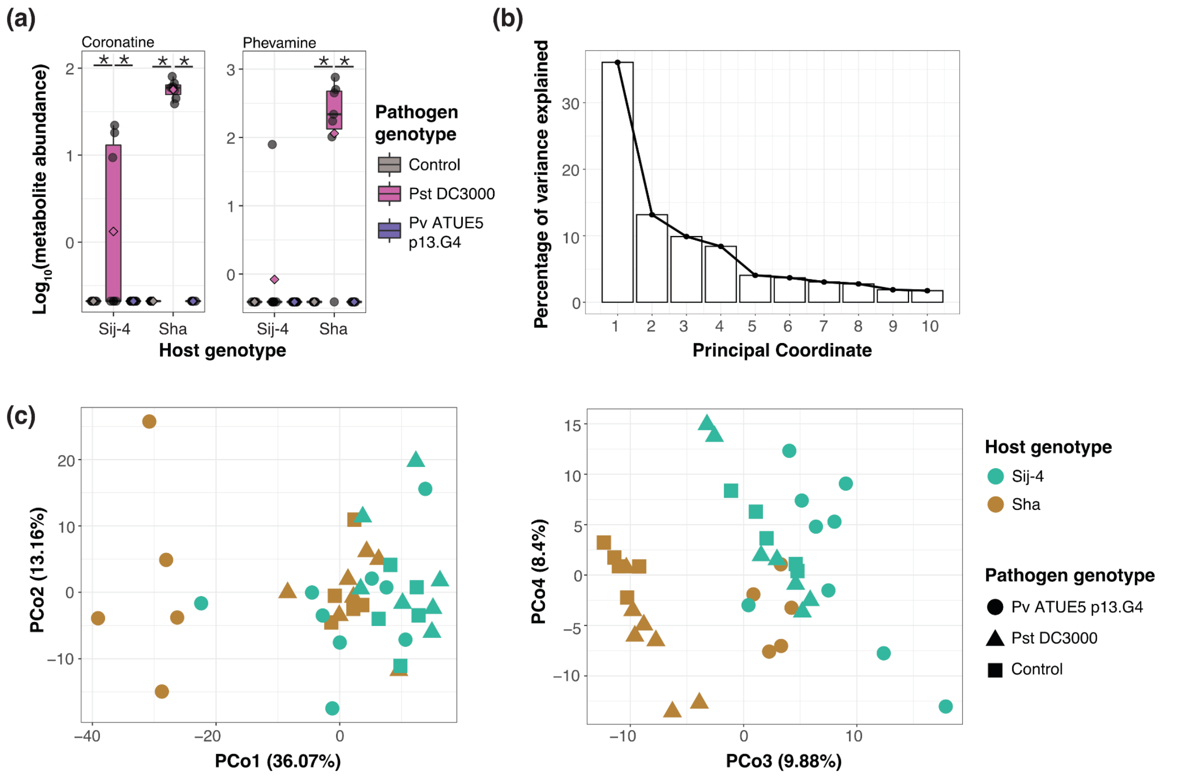


**Figure S9. Secondary metabolites profile of Sij-4 and Sha infected with Pv ATUE5** p13.G4 **and Pst DC3000. (a)** Coronatine and phevamine levels. (b) Scree plot of the Euclidean distance between samples based on 534 secondary metabolites. (c) Principal coordinate analysis of the Euclidean distance between samples based on 534 secondary metabolites. Colors indicate host genotypes and shapes indicate pathogen genotype. n = 5-9 replicates per host x pathogen genotype combination. Each replicate was a pool of 3 to 5 individual plants. Asterisks indicate statistically significant differences between two groups (Dunn's test, adj. p-value < 0.05).

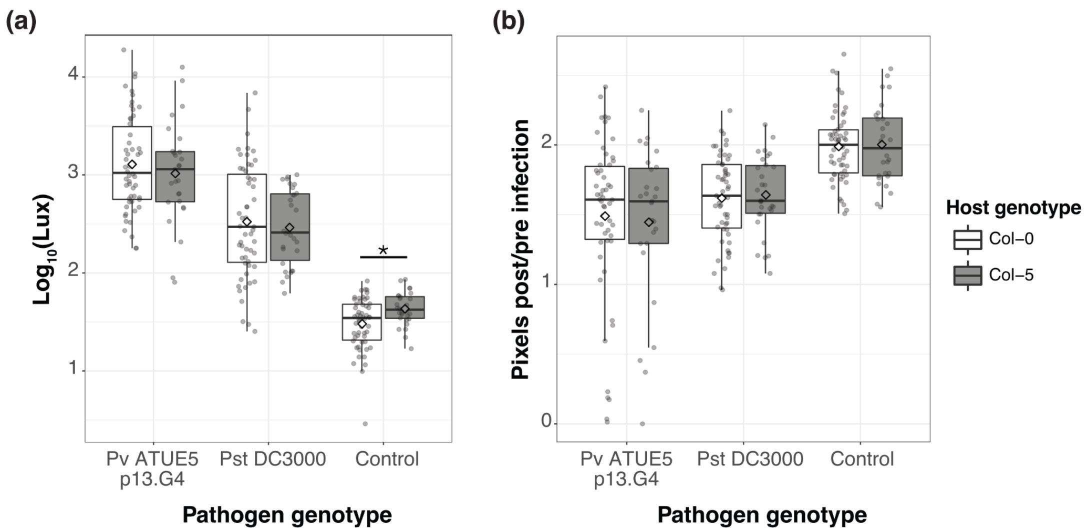


**Fig. S10. Col-0 and Col-5 are similarly susceptible to Pv ATUE5 and Pst DC3000. (a)** Bacterial load measured as luminescence 3 days post-infection. **(b)** Plant growth measured as the ratio of green pixels 3 days post-/pre-infection. Asterisks indicate statistical significance of the Wilcoxon test at p-value < 0.05 adjusted for multiple comparisons using the Benjamini-Hochberg method in comparison with Col-0 wild type within each pathogen genotype. n = 27-55 plants per host x pathogen genotype combination.

### Methods S1

#### Plant growth conditions

Seeds were sterilized by overnight incubation at -80 °C followed by at least 4 hours of vapor-phase sterilization with chlorine gas. Seeds were sown on Petri dishes with ½ MS medium with vitamins and MES buffer (Duchefa M0255.0050) and stratified for seven days at 4°C in the dark. Seedlings were grown under long-day (16 h) at 23°C in a growth chamber (Percival, model CU-36L5). After 3-5 days, seedlings were transferred to 24-well plates with the same medium, one seedling per well. 11-day-old plants were infected with single bacterial isolates.

#### Genetic distance

To calculate the genetic distance between *A. thaliana* accessions, single nucleotide polymorphisms (SNP) and indels were obtained from the data generated by the 1001 Genomes project data (1001Genomes, no date; 1001 Genomes Consortium, 2016). Host accessions Ey15-2, Koch-1 and Toufl-1 were not present in this dataset. Genetic distance was calculated using PLINK (Purcell *et al.*, 2007; Purcell, 2017) and plotted in R (RStudio Team, 2019; R Core Team, 2020).

#### Axenic infection

*Pseudomonas* isolates transformed with a *luxABCDE* reporter construct (Choi and Schweizer, 2006) were grown overnight at 28°C in Luria-Bertani (LB) medium with 100 µg/mL of nitrofurantoin (Sigma), diluted the following morning 1:10 in 5 mL selective medium and grown for 3 - 4 additional hours. Bacteria were pelleted at 3,500 g and brought to an $OD_{600}$ of 0.01 in 10 mM $MgSO_4$. 100 µL of this bacterial suspension were used to drip-inoculate 11-day-old plants on 24-well plates, distributing the volume over the whole rosette. Plants were mock infected with 10 mM $MgSO_4$ as control. Plates with plants were returned to the growth chamber, and whole rosettes were cut for analysis between 0- and 14-days post-infection.

Statistical analyses were performed using R v4.0.2 (R Core Team, 2020) and Rstudio (RStudio Team, 2019). ANOVA was used to identify significant differences between host genotype, bacterial genotype, and to determine the proportion of variance explained by each of these factors and their interaction. The model used for the ANOVA was $log_{10}$(luminescence) ~ pathogen genotype * host genotype + day + edge; where day corresponds to the day of the experiment and edge to whether the plant was on the outer wells of the 24-well plate. Variance decomposition was performed by dividing the ANOVA sum of squares of each variable by the total sum of squares. Significant differences among host or pathogen genotypes were determined using Tukey's Honestly Significant Differences (HSD) implemented in the function HSD.test of the *agricolae* package v1.3-5 (de Mendiburu, 2021). An example of the commands used:

```
agricolae::HSD.test(final.lm, "host_genotype", group = T, unbalanced = F.
```

Green pixels were quantified as described (Karasov *et al.*, 2020). Plates with plants were photographed with a tripod-mounted Lumix DMC-TZ61 digital camera. Plates were illuminated from below to prevent light reflection on the lid. Individual plants were extracted from whole-plate images by applying thresholds in Lab color space, followed by a series of morphological operations to remove noise and non-plant objects. Finally, a GrabCut-based postprocessing was applied and .csv files with plant IDs and green pixel counts were created. The workflow was implemented in Python 3.6 and *bash* using OpenCV 3.1.0 and *scikit-image* 0.13.0 for image processing. The ratio of green pixels was calculated by dividing the count of green pixels at a given time after infection by the count of green pixels before infection.

#### Segregation analysis

$F_2$ individuals derived from a Sha x Sij-4 cross (Chae *et al.*, 2014), Sij-4, and Sha individuals were grown and infected as described above. Plant size was extracted from pictures taken before and 3 days post-infection. We classified plants as resistant or susceptible based on the minimum green pixel ratio 3 dpi to 0 dpi of mock-infected $F_2$ plants. Statistical analyses were performed in R v4.0.2 (RStudio Team, 2019; R Core Team,

2020). Plant size before infection and green pixels ratio 3 days post-infection were compared using a linear regression pixels ~ host genotype, with experiment as a covariate when indicated. The binomial proportion confidence interval at 95% for the proportion of resistant plants was estimated using Wilson method as implemented in the package binom (Dorai-Raj, 2022). The experimental segregation ratio was compared to that expected from a 3:1 segregation using the Chi-squared goodness-of-fit test (chisq.test) implemented in the stats package from base R (R Core Team, 2020).

#### Transcriptomics

RNA extraction, mRNA library preparation and sequencing were performed as previously described (Yaffe *et al.*, 2012; Cambiagno *et al.*, 2021). Seven samples with more than 13 million reads were subsampled to obtain 13 million reads before analysis. We removed all samples with fewer than 3.4 million mapped reads from further analysis. One read was added to all read counts to avoid plotting -INF values in genes with read count 0 ($\log_{10}(0 + 1) = 0$; (Barragan *et al.*, 2021). Genes with less than ten counts over all samples were removed from downstream analyses. For exploratory data analysis, variance stabilizing transformed data were used and are referred here as normalized transcript counts. A gene was called as differentially expressed between two conditions when $|\log_2\text{FoldChange}| > 1$ and adj. p-value value < 0.01. Plots were generated using the R package ggplot2 (Wickham, 2016). Gene ontology enrichment analysis was performed with ShinyGO v0.76, available at http://bioinformatics.sdstate.edu/go/ (Ge, Jung and Yao, 2020). A list of differentially expressed genes was used as input for enrichment, and the 25,324 genes identified in this experiment were used as reference background. *Arabidopsis thaliana* was selected as species and 'GO biological process' as pathway database. Minimum (2) and maximum (2,000) pathway size were left as default, and the remove redundancy option was selected.

#### Metabolomics measurement and data analysis

Liquid chromatography-mass spectrometry (LC-MS) measurement of secondary metabolites (lipids, metabolites and dipeptides): the dried aqueous phase was measured using ultra-performance liquid chromatography (UPLC) coupled to a Q-Exactive mass spectrometer (Thermo Fisher Scientific) in positive and negative ionization modes, as described (Giavalisco *et al.*, 2011). Expressionist Refiner MS 12.0 (Genedata AG) was used for processing the LC-MS data with the following settings: chromatogram alignment (RT search interval 0.5 min), peak detection (minimum peak size 0.03 min, gap/peak ratio 50%, smoothing window 5 points, center computation by intensity-weighted method with intensity threshold at 70%, boundary determination using inflection points), isotope clustering (RT tolerance at 0.02 min, m/z tolerance 5 ppm, allowed charges 1 - 4), filtering for a single peak not assigned to an isotope cluster, charge and adduct grouping (RT tolerance 0.02 min, m/z tolerance 5 ppm). An in-house library of authentic reference compounds was used to identify molecular features allowing 10 ppm mass deviation and dynamic retention time deviation (maximum 0.15 min).

Ultra-performance liquid chromatography electrospray ionization tandem mass spectrometry (UPLC-ESI-MS/MS) measurement of hormones: a fixed volume (0.250 ml) of upper supernatant (MTBE phase) was transferred to a fresh 1.5-ml microcentrifuge tube and dried down using a SpeedVac concentrator at 25. Dried pellets were resuspended in 100 µl of water:methanol (50:50) solution and the resuspended samples were immediately subjected to UPLC-ESI-MS/MS analysis. Hormones were measured as described (Salem *et al.*, 2020)) using a quadrupole/linear ion trap (QLIT) mass analyzer (4000 QTRAP; AB Sciex Germany) with a multiple-reaction monitoring (MRM) scan type equipped with an electrospray ionization (ESI) source (Turbo Ion Source; AB Sciex) and attached to the UPLC system. *analyst* 1.6.2 (AB Sciex) was used for instrument control, data acquisition, processing, and analysis (Salem *et al.*, 2020).

Data analysis: median-normalized secondary metabolite and hormone abundances were $\log_{10}$-transformed, scaled and centered before further analysis. Metabolites that did not vary among samples were removed. Statistical tests and plotting were performed using R (R Core Team, 2020) and Rstudio (RStudio Team, 2019). To detect changes between treatments, principal coordinate analysis (PCoA) was performed using the *cmdscale* function. Permutational multivariate analysis of variance (PERMANOVA) was conducted based on Euclidean distance between samples, using the function *adonis2* of the *vegan* package

(Oksanen *et al.*, 2020). To detect statistically significant differences in metabolite abundances between treatments within each host genotype, a Kruskal-Wallis test was performed, followed by Dunn's test with Benjamini-Hochberg correction for multiple comparisons, as implemented in the dunn.test package (Dinno, 2017).
